# Endothelial LRRC8C associates with LRRC8A and LRRC8B to regulate vascular reactivity and blood pressure

**DOI:** 10.1101/2025.08.17.670763

**Authors:** Qiujun Yu, Yonghui Zhao, Joshua Maurer, Prakash Arullampalam, Nathaniel John, John D. Tranter, Tarek Mohamed Abd El-Aziz, Michelle Lin, Daniel J. Lerner, Carmen M. Halabi, Rajan Sah

## Abstract

Vascular tone is impacted by the endothelium’s ability to detect mechanical and chemical stimulation. *L*eucine-*R*ich *R*epeat-*C*ontaining protein *8*A, (LRRC8A), was previously identified as a required component of the mechanoresponsive endothelial LRRC8 complex regulating AKT-endothelial nitric oxide synthase (eNOS) signaling and vascular function. While LRRC8A is broadly expressed, LRRC8B, C, D and E have tissue-restricted expression. Here, we identified 2 single nucleotide polymorphisms (SNPs) in *LRRC8C* highly associated with elevated diastolic and systolic blood pressure in human genetic studies, implicating LRRC8C as a regulator of vascular function. While LRRC8A/B/C/D/E are expressed in endothelium, co-immunoprecipitation experiments from lung endothelium using *Lrrc8a*-3xFlag knock-in mice, *Lrrc8c-*HA knock-in mice and endothelium-specific *Lrrc8a*-3xFlag overexpression mice demonstrate the endothelial LRRC8 complex to be composed largely of LRRC8A/B/C heteromers. *Lrrc8a/b/c* knock-out studies in mice and knock-down studies in human umbilical vein endothelial cells show co-dependent expression of LRRC8A/B/C proteins, but not LRRC8D. Functionally, LRRC8A and LRRC8C depletion reduces endothelial volume regulatory anion channel (VRAC) currents, inhibits AKT-eNOS signaling, increases myogenic tone, impairs eNOS dependent vasodilation, and exacerbates angiotensin-induced hypertension. These data identify LRRC8A, LRRC8B and LRRC8C as components of the endothelial LRRC8 complex and reveal LRRC8C as having a non-redundant role in regulating endothelial AKT-eNOS, vascular relaxation and susceptibility to hypertension.

## Introduction

Vascular tone is impacted by the endothelium’s ability to detect both mechanical and chemical stimulation. Endothelial Akt-nitric oxide synthase (eNOS) signaling is known to relax vascular smooth muscle cells via endothelial nitric oxide production, causing vasodilation and reducing vascular resistance, thereby regulates blood pressure and organ perfusion. Small resistance arteries and arterioles have intrinsic mechanisms to regulate blood flow that depend on mechanical factors, such as shear stress exerted by blood flow or wall stress exerted by intraluminal blood pressure. The latter mechanism, termed myogenic response or Bayliss effect, is characterized by vasoconstriction induced by increasing intraluminal pressure ^1^. While myogenic response in resistance arteries contributes to vascular autoregulation which maintains capillary hydrostatic pressure and blood flow over a wide pressure range ^2^, persistent elevation of myogenic tone increases vascular resistance and remodeling, enhances vascular sensitivity to vasoconstrictors and ultimately leads to development and progression of hypertension.

The endothelial volume regulatory anion channel (VRAC) has been proposed to be mechano-sensitive and to activate in response to fluid flow/hydrostatic pressure and putatively regulate vascular reactivity. We recently reported that the Leucine Rich Repeat Containing Protein 8A (LRRC8A) is a required component of the heterohexameric complex that forms VRAC in human umbilical vein endothelial cells (HUVECs) ^3^. Endothelial LRRC8A associates with GRB2, regulates eNOS, AKT and mTOR signaling under basal conditions, with stretch and shear-flow stimulation and is required for endothelial cell (EC) alignment to laminar shear flow. Endothelium-restricted *Lrrc8a* KO (e*Lrrc8a* KO) mice develop hypertension in response to chronic angiotensin II infusion and exhibit impaired retinal blood flow with both diffuse and focal blood vessel narrowing in the setting of diabetes. Having established the significance of LRRC8A in regulating vascular function, there remains a knowledge gap in the molecular identity of specific LRRC8 heteromers that form VRAC in endothelium.

The LRRC8 complex is comprised of heterohexamers of LRRC8A and combinations of LRRC8B, C, D or E, where LRRC8A is the essential subunit to traffic other LRRC8 subunits to the plasma membrane and form viable channels ^4,5^. While LRRC8A is broadly expressed, LRRC8B, C, D and E have tissue-restricted expression patterns. Due to the combinatorial nature of LRRC8 biology the role of LRRC8B-E as the function-refining subunits vary in different tissues ^6^. Importantly, differences in LRRC8 subunits composition regulates LRRC8 channel permeability and conductivity ^5,7,8^. Therefore, defining the specific LRRC8 protein complex in endothelial cells is necessary to understand the molecular basis of LRRC8 signaling in endothelium and to potentially also guide tissue-targeted drug design.

In this study, we identify genetic variations in *LRRC8C* which reveals strong association with diastolic and systolic blood pressure. We show that endothelial LRRC8 complex is composed largely of LRRC8A/B/C heteromers which possess a co-dependent expression pattern. More importantly, we demonstrate LRRC8C has a non-redundant role in contributing to endothelial VRAC currents, regulating endothelial AKT-eNOS signaling, vascular relaxation and susceptibility to hypertension – supporting the association identified from human genetic studies.

## Methods

### Sex as a biological variable

Male and female mice were included in all experimental procedures, and the results demonstrated no statistically significant differences between sexes. Therefore, sex was not considered a biological variable in the study. All aspects of the experimental design, data acquisition, and statistical analysis were conducted without stratification by sex.

### Animals

All animal-related experimental procedures were conducted in accordance with ethical guidelines and received approval from the Institutional Animal Care and Use Committee (IACUC) at Washington University School of Medicine. *Lrrc8a^fl^*^/fl^ (Swell1^fl/fl^) mice were generated as described previously ^9,10^. *CDH5-Cre;Lrrc8a^fl/fl^* (endothelial specific Lrrc8a knockout, *eLrrc8a* KO) mice were generated as described previously ^3^. *Lrrc8a*-3xFlag knock-in (3xFlag KI) mice were generated by inserting a 3xFlag epitope tag into the endogenous C-terminal locus of the *Lrrc8a* gene. Endothelial specific *Lrrc8a*-3xFlag overexpression (e*Lrrc8a* OE) mice were generated by crossing CAG-LSL-*Lrrc8a*-3xFlag transgenic mice (generated by Cyagen Biosciences, Santa Clara, CA) with CDH5-Cre mice. CDH5-Cre mediated excision of a floxed STOP codon releases endothelium-specific *Lrrc8a*-3xFlag expression. *Lrrc8c*-HA knock-in (*Lrrc8c*-HA KI) mice were generated by inserting a HA epitope tag into the endogenous *Lrrc8c* C-terminal locus. g*Lrrc8b* KO mice were generated by CRISPR/cas9-mediated gene-editing using the Genome Engineering & Stem Cell Center (GESC@MGI) at the Washington University in St. Louis. g*Lrrc8c* KO mice were a gift from Dr. Axel Concepcion (University of Chicago, Chicago, IL). All mice were maintained under standardized housing conditions, including controlled temperature, humidity, and a 12-hr light/dark cycle. Food and water were available ad libitum. Mice with ages ranging from 8 to 20 weeks were utilized in the study. Both *in vitro* and *in vivo* experiments were conducted using mixed gender with age and gender matched among experimental groups.

### Antibodies

Anti-LRRC8A antibody was developed as described previously ^9,10^. Anti-LRRC8B and anti-LRRC8E antibodies were purchased from Sigma (St. Louis, MO, t#ZRB2950 and #ZRB2957). Anti-LRRC8D was kindly provided by Dr. Thomas Jentsch (Leibniz-Forschungsinstitut für Molekulare Pharmakologie, Berlin, Germany) and anti-LRRC8C was a gift from Dr. Axel Concepcion (University of Chicago, Chicago, IL). All LRRC8 antibodies were validated in knockout and knockdown studies. All other primary antibodies were purchased from Cell Signaling (Danvers, MA): Akt1 (#2938), Akt2 (#3063), p-Akt1 (#4060), p-Akt2 (#8599), eNOS (#32027), p-eNOS (#9571), anti-,8-actin (#8457). Rabbit IgG was purchased from Santa Cruz (Dallas, TX, #sc-2027). Anti-CD31 was purchased from Thermo Fisher (Waltham, MA, #MA3105).

### Cell culture

Human umbilical vein endothelial cells (HUVECs) were purchased from ATCC (#PCS-100-013). Cells were grown on 1% gelatin-coated plates at 37°C with 5% CO_₂_ in a humidified incubator. The culture medium consisted M199 supplemented with 20% FBS, 0.1 mg/mL heparin sodium salt (Alfa Aesar, Ward Hill, MA, #9041-08-1), and 0.03 mg/mL ECGS (Millipore Sigma, t#02-102).

### Adenovirus infection in HUVECs

HUVECs were plated at 360,000 cell/well in six-well plates and were grown for 24 hr to reach 70% confluency. Cells were infected with either human adenovirus type 5 with sh*Lrrc8a*(Ad5-mCherry-U6-sh*Lrrc8a*-shRNA, 2.2 × 10^10^ PFU/ml, Vector Biolabs, Devault, PA, # shADV-214592), shLrrc8c(Ad5-mCherry-U6-h*Lrrc8c*-shRNA, 2.2 ×10^10^ PFU/ml, Vector Biolabs, # shADV-214595), or a scrambled non-targeting control (Ad5--mCherry-U6-scramble, 1 ×10^10^ PFU/ml) at a multiplicity of infection of 50 for 12 hr, and studies performed 3 - 4 days after adenoviral transduction. The targeting sequences are GCA CAA CAT CAA GTT CGA CGT for sh*Lrrc8a*, and CGA TTA CCT CTC AGT AGC CAT for sh*Lrrc8c*.

### SiRNA transduction in HUVECs

si*Lrrc8a* (ID# s32109), si*Lrrc8b* (ID# s23956), si*Lrrc8c* (ID# s38687), si*Lrrc8d* (ID# s106871), along with scramble control siRNA (# 4390843) were purchased from Invitrogen. HUVECs were seeded at a density of 360,000 cell/well in six-well plates and cultured for 24 hr to achieve approximately 70% confluency. Cells were transduced twice with silencer select siRNA at 24 and 72 hr post-plating. Each siRNA was combined with Opti-MEM (285.25 µl, #11058-021, Invitrogen, Waltham, MA), siPORT amine (8.75 µl, #AM4503, Invitrogen) and the silencer select siRNA (6 µl) in a final volume of 300 µl. HUVECs were transduced for 4 hr at 37°C in DMEM supplemented with 1% FBS. Following transduction, cells were returned to standard growth media consisting of M199, 20% FBS, 0.05 g heparin sodium salt, and 15 mg ECGS. Cell lysates were harvested on day 4 under basal conditions or after 16 hr of serum starvation in FBS-free culture media.

### Electrophysiology

Whole-cell recordings were performed at room temperature using Axopatch 200B amplifier or a MultiClamp 700B amplifier paired with a Digidata 1550 digitizer and both used pClamp 10.4 software (Molecular Devices, San Jose, CA). For standard VRAC whole-cell recordings, the intracellular solution contained (in mM): 120 L-aspartic acid, 20 CsCl, 1 MgCl2, 5 EGTA, 10 HEPES, 5 MgATP, 120 CsOH, 0.1 GTP, pH 7.2 with CsOH. The isotonic extracellular solution contained (in mM): 90 NaCl, 2 CsCl, 1 MgCl_2_, 1 CaCl_2_, 10 HEPES, 110 mannitol, pH 7.4 with NaOH (300 mOsm/kg). For hypotonic swelling (210 mOsm/kg), the extracellular solution was modified by replacing 110 mM mannitol with 10 mM mannitol, while maintaining all other components. Swell-activated VRAC currents were induced by perfusing cells with a hypotonic solution (210 mOsm/kg). Osmolarities were measured using a Wescor Vapro 5520 Osmometer (ELITechGroup Inc., Logan, UT). Patch pipettes, pulled from borosilicate glass capillaries (WPI) using a P-87 puller (Sutter Instruments, Novato, CA), exhibited a resistance of ∼4–6 MΩ when filled with intracellular solution. The holding potential was 0 mV. Voltage ramps from -100 to +100 mV (at 0.4 mV/ms) were applied every 4 s. Currents were filtered at 10 kHz and sampled at 100 ms interval.

### Immunoblotting

Mouse lung tissue was harvested after right ventricle injection of ice-cold phosphate buffered saline. Tissues were snap-frozen in liquid nitrogen and lysed in ice-cold lysis buffer (150 mM NaCl, 20 mM HEPES, 1% NP-40, 5 mM EDTA, pH 7.4) containing1:10 PhosSTOP™ Roche Diagnostics, Indianapolis, IN, #4906845001) and 1:20 cOmplete™ ULTRA Protease Inhibitor Cocktail (Roche Diagnostics, #06538304001). Tissue lysates were kept on ice with gentle agitation for 20 minutes to allow complete lysis. Lysates were cleared of debris by two cycles of centrifugation at 12,500 × g for 20 min at 4°C. Resulting supernatants were transferred to fresh tubes and solubilized protein was measured using a DC protein assay kit (#5000111, Bio-Rad, Hercules, CA). For immunoblotting, samples containing 10 µg of protein were mixed with 4x Laemmli sample loading buffer (Bio-Rad) containing 1:10 β-mercaptoethanol, heated at 90°C for 5 minutes, and then loaded onto a 4–20% Mini-PROTEAN® TGX™ Precast Protein Gels (#4561083, Bio-Rad) which were ran in tris/glycine/SDS running buffer (#1610772, Bio-Rad) at 100 V for 1.5 – 2h. After separation, gels were transferred to an immune-Blot PVDF membrane (#1620177, Bio-Rad) in tris/glycine buffer (#1610771, Bio-Rad) for at 100 V for 1.5 h at 4°C. Membranes were then blocked with either 5% bovine serum albumin (#A-420-10, GoldBio, St. Louis, MO), or milk, in tris-buffered saline with Tween (TBS-T; 0.2 M Tris, 1.37 M NaCl, 0.2% Tween-20, pH 7.4) at room temperature for 30 minutes. Membranes were incubated with primary antibody at a concentration of 1:1000 in 1% BSA or milk overnight at 4°C. Membranes were washed three times in TBS-T prior to incubation with secondary antibodies (Goat-anti-mouse #170-5047, Goat-anti-rabbit #170-6515, Bio-Rad) at a concentration of 1:5000 in 1% BSA or milk at room temperature for 1 hr. After washing three times, membranes were incubated with Clarity™ Western ECL substrate (#1705060, Bio-Rad) and visualized using the ChemiDoc Imaging system (Bio-Rad). Protein band intensities were quantified using ImageJ, with β-actin used as a loading control.

### Immunoprecipitation

Lung tissue lysates from both *Lrrc8a*-3xFlag KI, e*Lrrc8a* OE and *Lrrc8c*-HA KI mice were incubated with anti-Flag and anti-HA monoclonal antibody-bound magnetic beads (#M8823, Sigma, and #88837, Pierce) overnight at 4°C under continuous end-over-end rotation. The anti-Flag bound magnetic beads were then separated on magnetic stand and washed 3 times with RIPA buffer. Bound proteins were eluted in Laemmli buffer (Bio-Rad), boiled at 95°C for 5 minutes, and subsequently run on SDS-PAGE followed by Western blot analysis.

### Measurement of myogenic tone in mesenteric arteries

Following euthanasia with CO_2_, the gut from the duodenum to the cecum was excised and placed into cold normal Krebs’ buffer containing (in mM): 120 NaCl, 25 NaHCO_₃_, 4.8 KCl, 1.2 NaH_₂_PO_₄_, 1.2 MgSO_₄_, 11 glucose, and 1.8 CaCl_₂_ (pH 7.4). The third-order mesenteric arteries were isolated, mounted, and secured using nylon sutures in a pressure myography system (#114P, Danish Myo Technology, Denmark), as previously described ^11^. For the endothelial denuded experiments, air bubbles were passed through the mesenteric artery to remove the endothelial cell layer. To ensure there were no leaks, the vessel was pressurized to a physiological pressure of 60 mmHg for less than one minute.

Endothelial removal was confirmed by observing the loss of vasodilation to 10 μM Acetylcholine at 60 mmHg in 10 μM phenylephrine (PE)-pre-constricted arteries. The pressure was then lowered to 10 mmHg to allow vessels to equilibrate for 30 minutes at 37°C. Vessels then underwent 10-minute pressure steps ranging from 10 mmHg to 120 mmHg, to allow for the development of stable myogenic tone. After completing the calcium-dependent assay, the vessels were exposed to a calcium-free Krebs’ buffer (same composition but added with 2 mM EGTA). Data during each pressure step were recorded at the end of the 10-minute pressure step and expressed as a percentage of the diameter change relative to the initial diameter [(D_x_ – D_10_)/D_10_*100]. Myogenic tone is calculated by measuring relative difference in active and passive diameters and expressed as (passive diameter – active diameter)/passive diameter x 100%.

### Measurement of vasodilatory reactivity in mesenteric arteries

Mesenteric arteries were isolated and mounted in the pressure myography system as above. Arteries were equilibrated at 37°C for 30 min under a pressure of 10 mmHg, with initial responsiveness tested using 10 μM phenylephrine (PE) and 10 μM acetylcholine chloride (Ach) at 60 mmHg. Vessels showing at least 40% dilation were selected for subsequent experiments. Vessels were pre-constricted with 10μM PE and outer diameter measurements were taken after 1 min using increasing concentrations of Ach (1 nM-1 mM) or papaverine (1 nM-1 mM), recorded with the MyoVIEW 4 software. In experiments involving nitric oxide synthase (NOS) inhibition, L-N^G^-Nitro arginine methyl ester (L-NAME, 10μM) was added to the bath superfusate and vessels were incubated at 10 mmHg for 30 min prior to testing Ach and papaverine responses in the presence of L-NAME. The average response of 3 vessels per agonist was calculated for each mouse, with 3-4 mice per genotype. All reagents including phenylephrine, acetylcholine chloride, papaverine and L-NAME were purchased from Sigma.

### Angiotensin II infusion

Alzet osmotic minipumps (Model 1004) were used to deliver angiotensin II (Bachem, Torrance, CA), dissolved in saline, at a rate of 600 ng/kg/min for 4 weeks. Under 2% isoflurane anesthesia, mice underwent subcutaneous implantation of the minipumps via a ∼1 cm incision behind the ear over the shoulder blade of the front leg. 1-2 wound clips were placed to close the incision, and topical lidocaine was applied to the wound.

### Non-invasive blood pressure measurement

Tail-cuff blood pressure (BP) measurements were performed at a consistent time each dayusing computerized tail-cuff system BP-2000 (Visitech Systems, Allen, Texas). Mice were acclimated to the tail-cuff system by performing 3 consecutive days of measurements with 20 sequential measurements each day. Baseline BP was calculated as the average of measurements taken over 3 days, excluding the acclimation period. BP was then measured weekly for 4 weeks, beginning 7 days after angiotensin II minipump implantation. At the end of the 4-week period, post-implantation BP was determined by averaging 3 consecutive days of measurements.

### Statistics

Data are represented as mean ± SEM. Two-tail unpaired Student’s t-tests were used for comparison between two groups. For three or more groups, data were analyzed by one-way analysis of variance (ANOVA) and Tukey’s post hoc test. For time-series data between two groups, data were analyzed using a two-way ANOVA followed by Bonferroni post hoc test. A probability of *p* < 0.05 was considered to be statistically significant.

### Study Approval

All experimental procedures involving mice were approved by the Institutional Animal Care and Use Committee (IACUC) of Washington University in St. Louis.

### Data availability

Supporting data values associated with the main manuscript and supplement materials are provided in the Supporting Data Values file.

### Author contributions

QY, YZ, JM, and RS designed research studies. QY, YZ, JM and RS conducted experiments, analyzed data, and prepared figures. PA assisted with patch clamp study. NJ assisted with immunoprecipitation study. ML and CH performed pressure myography experiments in mesenteric arteries. DL analyzed and interpretated the GWAS and PheWAS data. QY and RS wrote the manuscript. JM, JDT, TMAEA, ML and CH revised the manuscript. All authors reviewed and approved the manuscript.

## Results

### Genome-wide Association Studies identified two *LRRC8C* SNPs associated with systolic and diastolic blood pressure

We recently demonstrated that *Lrrc8a* functionally encodes endothelial VRAC current and regulates endothelial response to mechanical stretch and shear flow via PI3K/AKT/eNOS and ERK signaling ^3^. In addition, endothelial targeted Lrrc8a ablation in vivo predisposes to angiotensin-II induced hypertension and microvascular dysfunction in the setting of high-fat high-sucrose diet induced diabetes ^3^. Based on these findings, we hypothesized that LRRC8 channels may be important in human blood pressure regulation. In humans, the five LRRC8 subunits are encoded on three chromosomes (LRRC8A 9q34.11, LRRC8B-D 1p22.2, LRRC8E 19p13.2) 1. We searched the National Health Genome Research Institute-European Bioinformatics Institute (NHGRI-EBI) catalog of Genome-wide Association Studies (GWAS) (ebi.ac.uk/gwas/home) and identified 2 exonic SNPs, rs12032393 and rs113563325 in *LRRC8C* that are significantly associated with diastolic and systolic blood pressure (BP), respectively (**Fig. 1A**).

**Figure 1.**
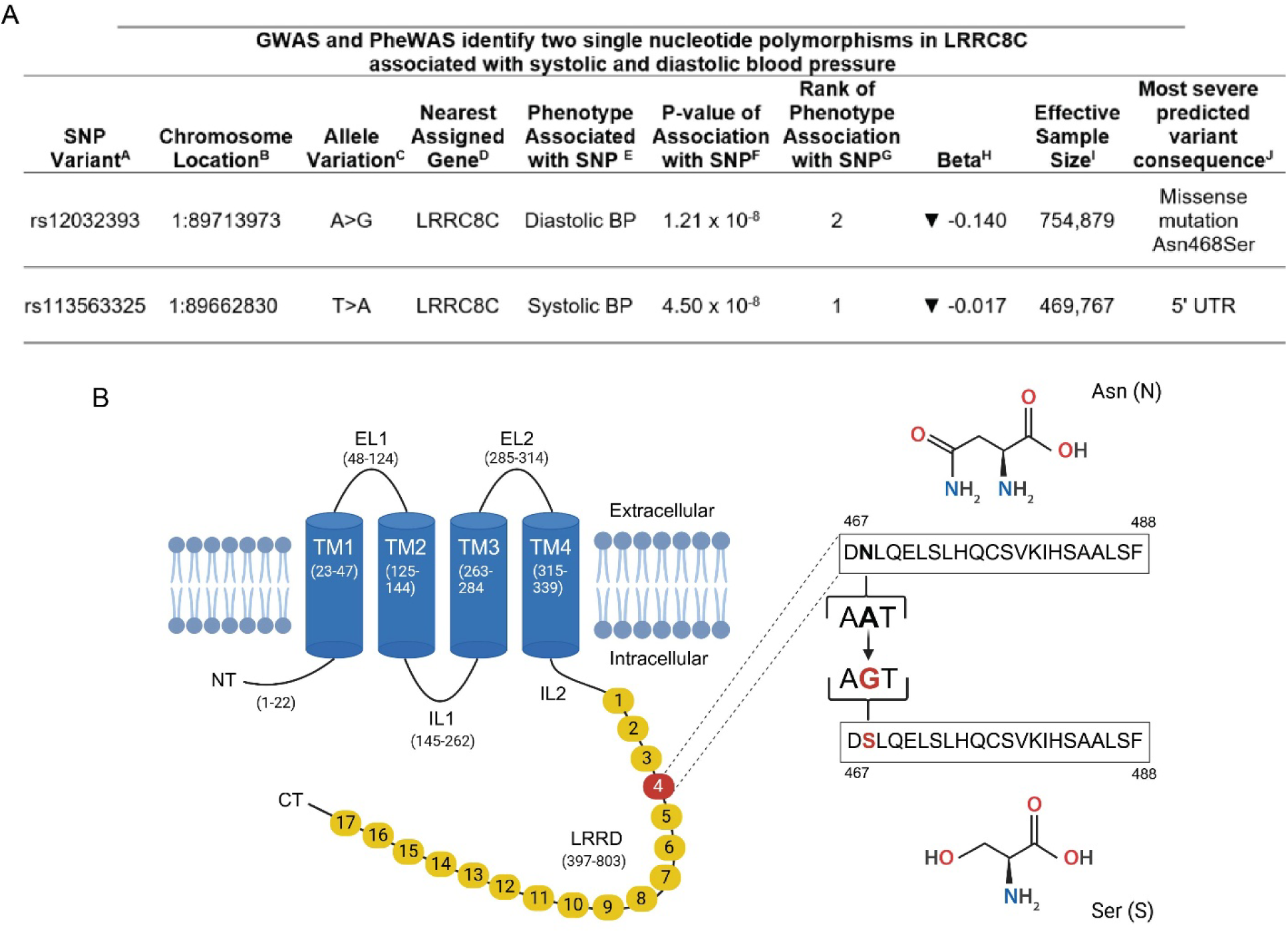
Single nucleotide polymorphisms (SNP) in Leucine-Rich Repeat-Containing protein 8C (*LRRC8C*) associates with blood pressure in human genetic studies. (**A**) GWAS and PheWAS identify two SNPs in *LRRC8C* associated with systolic and diastolic blood pressure. ^A^SNP variants associated with BP in the GWAS Catalog (https://www.ebi.ac.uk/gwas/home). ^B^GRCh38 position (chromosome:nucleotide) from Single Nucleotide Polymorphism database (dbSNP) (https://www.ncbi.nlm.nih.gov/snp/). ^C^Common allele > variant allele. ^D^Nearest gene predicted by Ensembl Variant Effect Predictor (https://useast.ensembl.org/info/docs/tools/vep/index.html). ^E^BP phenotype associated with SNP; ^F^P-value of the association of the BP phenotype with the SNP (IEU Open GWAS project (https://gwas.mrcieu.ac.uk/)). ^G^Rank of the association of the BP phenotype with the SNP compared to other phenotypes associated with the SNP, from most (1) to least significant. ^H^Direction and per unit change associated with the SNP variant. ^I^Number of people that contributed alleles to the study; ^J^Most severe variant consequence predicted using Ensembl Variant Effect Predictor (https://useast.ensembl.org/Multi/Tools/VEP). **(B)** A schematic representation of the A>G mutation at SNP rs12032393 replacing 468 asparagine to serine. 5’ UTR, 5-prime untranslated region; BP, blood pressure; CT, C-terminus; EL, extracellular loop; GWAS, Genome-Wide Association Study; IL, intracellular loop; LRRD, Leucine-Rich Repeat-Containing Domain; NT, N-terminus; PheWAS, Phenome-Wide Association Study; TM, transmembrane domain.

The two SNPs are not in linkage disequilibrium in the 1000 Genome population or the 5 sub-populations (r2 0.0052, D’ 0.485) (https://ldlink.nih.gov/?tab=ldpop). One of these (rs12032393) is an exonic A->G variant that confers a missense mutation replacing asparagine 468 with serine. This mutation is located in the 4th leucine-rich repeat region of the 803 amino acid protein, converting a polar amide side chain on (N) to a less polar hydroxyl side chain in (S) (**Fig. 1B**). This variant allele is associated with a decrease in diastolic BP. The other SNP (rs113563325) is in 5’ UTR3 and is associated with a decrease in systolic BP. Data from publicly available Phenome-wide Association Studies (PheWAS) reveal blood pressure to be the first-or second-strongest association with the two *LRRC8C* SNP variants (**Fig. 1A**). The beta value, or impact, of the missense SNP is almost 10-fold that for the 5’UTR variant (−0.14 vs −0.017), suggesting the missense SNP has substantially more impact on diastolic BP than the 5’UTR variant on systolic BP. The frequencies of the missense SNP risk allele A>G (24%) and the G|G homozygous genotype (5.5%) are similar across regional populations, except for the population with African ancestry which has dramatically lower frequency of the risk allele (A>G (3%) than all other regions (https://useast.ensembl.org/Homo_sapiens/Variation/Population?db=core;r=1:89713473-89714473;second_variant_name=rs2065152;v=rs12032393;vdb=variation;vf=548616). The frequency of the G|G genotype was not provided. While the frequencies of the 5’UTR SNP risk allele T>A (1.4%) and the A|A homozygous genotype (0.2%) are also lowest in the population of African ancestry, the distribution in the other population regions is more variable, 6.5% - 26.9% and 1.2% v. 1.1 to 8.0% than for the missense SNP. (https://useast.ensembl.org/Homo_sapiens/Variation/Population?db=core;r=1:89662330-89663330;v=rs113563325;vdb=variation;vf=1646593). Therefore, there is a clear human genetic signal for the importance of the LRRC8 complex in blood pressure regulation, and specifically LRRC8C.

### Lung tissue is highly enriched in endothelial LRRC8 proteins

Lung is the most vascularized organ and endothelial cells in the pulmonary vascular bed account for the majority of cells in the lungs. Therefore, we turned to lung tissue to study endothelial LRRC8 signaling in vivo ^12^. As shown in **Figure 2 A-D**, both VEGF (**Fig. 2A, C**) and LRRC8A (**Fig. 2A-B**) are highly expressed in wildtype (*Lrrc8a fl/fl*) lung tissue lysates, and ∼90% of this LRRC8A protein is absent in endothelial cell (EC) specific *Lrrc8a* knockout mice (CDH5-Cre;*Lrrc8a fl/fl,* eLRRC8A KO, **Fig. 1A,B,D**), suggesting the majority of the LRRC8A signal from lung is EC derived. Indeed, the same pattern of VEGF and LRRC8A is observed in the mesenteric arterial beds from *Lrrc8a fl/fl* and e*Lrrc8a* KO mice (**Fig. 1E-H**), further supporting the use of lung tissue as a surrogate for studying LRRC8 composition and signaling in primary endothelium in mice.

**Figure 2.**
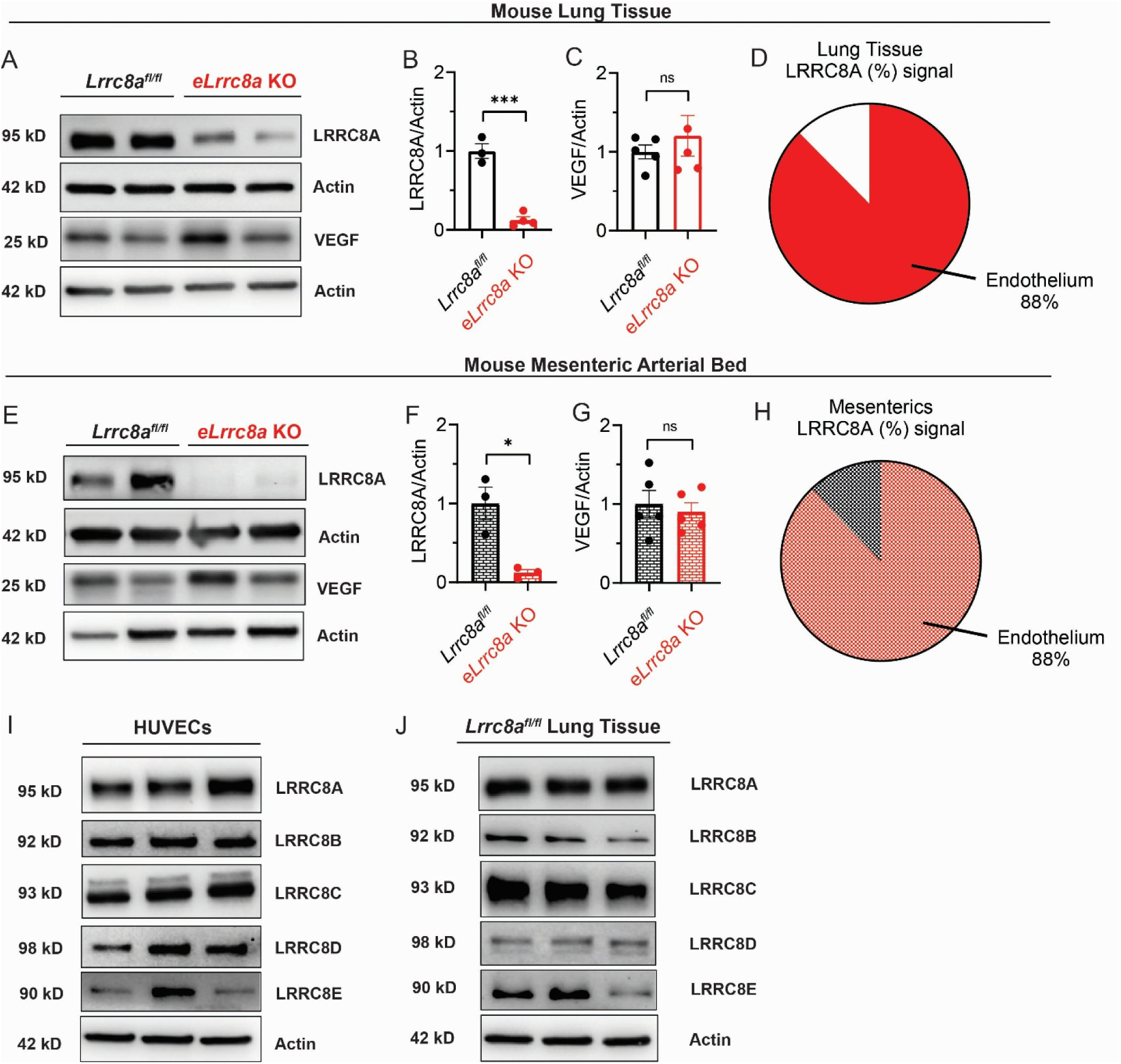
Leucine-Rich Repeat-Containing protein 8 (LRRC8) subunits are highly expressed in murine pulmonary endothelium. **(A)** LRRC8A, vascular endothelial growth factor (VEGF), and β-actin protein expression in lung tissues from endothelial specific *Lrrc8a* knockout (*eLrrc8a* KO) and *Lrrc8a*^flfl^ mice. **(B, C)** Densitometric quantification of LRRC8A/β-actin (n = 3-4) and VEGF/β-actin (n = 4-5). **(D)** Pie chart showing an 88% reduction of LRRC8A protein levels in lung tissue from *eLrrc8a* KO mice as compared to *Lrrc8a^flfl^*control mice. **(E, G)** LRRC8A (n = 3), VEGF (n = 5) and β-actin protein expression in meserteric arterial bed from *eLrrc8a* KO and *Lrrc8a^flfl^* mice (**H**) Pie chart showing an 88% expression reduction of LRRC8A protein levels in the mesenteric arterial bed from *eLrrc8a* KO mice as compared to *Lrrc8a^flfl^* control mice. **(I, J)** LRRC8A, LRRC8B, LRRC8C, LRRC8D, and LRRC8E protein expression in human umbilical vein endothelial cells (HUVECs) and *Lrrc8a^flfl^* mouse lung tissue. Statistical significance between indicated groups was calculated using a two-tailed unpaired Student’s t-test. Data are represented as mean ± S.E.M. ns - not significant; \**p* < 0.05; \*\*\**p* < 0.001.

### Endothelial LRRC8 complex is a LRRC8A/B/C heteromer

Since the LRRC8 complex is composed of different combinations of LRRC8A-E, with potentially different functions, we profiled LRRC8A-E protein expression in human umbilical vein endothelial cells (HUVEC, **Fig. 2I**) and mouse lung lysates (**Fig. 2J**) by western blot. All LRRC8 subunits are expressed in endothelial cells in vitro and in vivo (**Fig. 2I-J**). We next sought to determine the specific LRRC8 complex in primary endothelium by co-immunoprecipitation (IP) using lung lysates. As we found no antibodies capable of specific LRRC8A or LRRC8C IP, we generated 3 mouse models wherein 3XFlag epitopes were fused to the *Lrrc8a* C-terminus to allow for co-IP using more specific Flag antibodies: 1. *Lrrc8a*-3xFlag knock-in (*Lrrc8a*-3xFlag KI) mice in which a 3xFlag epitope was knocked into the endogenous *Lrrc8a* C-terminal locus using CRISPR/Cas9 gene-editing (**Fig. 3A, C, D**); 2. Endothelium-specific *Lrrc8a*-3xFlag overexpression (e*Lrrc8a*-3XFlag OE) mice wherein CDH5-Cre-mediated excision of a floxed STOP codon in a CAG-LSL-*Lrrc8a*-3xFlag transgene induces mild endothelium-specific *Lrrc8a*-3xFlag over-expression (**Fig. 3B, E, F**); 3. *Lrrc8c*-HA knock-in (*Lrrc8c*-HA KI) mice in which a HA epitope was knocked into the endogenous *Lrrc8c* C-terminal locus using CRISPR/Cas9 gene-editing (**Fig. S1A**). Immunoprecipitation with 3xFlag antibody yielded efficient LRRC8A pulldown from lung lysates isolated from *Lrrc8a-*3xFlag KI mice, and further western blot of the IP complex revealed presence of LRRC8B, C, D and E subunits in LRRC8A-containing heteromers from lung lysates of *Lrrc8a*-3xFlag KI mice, with a strong signal for LRRC8C (**Fig. 3G**). Similar results were found using lung lysates of *Lrrc8c*-HA KI mice where LRRC8C pulldown by HA antibody IP followed by western blot demonstrated presence of all LRRC8 subunits **(Fig.S1B)**. However, IP-Western blot from e*Lrrc8a*-3xFLAG lung lysates favored primarily LRRC8A/B/C heteromer, again with prominent LRRC8C signal (**Fig. 3H**). As IP of LRRC8 complexes using e*Lrrc8a*-3XFlag OE lung lysates specifically targets endothelium as opposed to other cell-types present in lung tissues, these data suggest that the endothelial LRRC8 complex is primarily an LRRC8A/B/C heteromer, while the LRRC8D and LRRC8E contributions observed in *Lrrc8a*-3xFlag KI and of *Lrrc8c*-HA KI likely arise from non-endothelial cell types.

**Figure 3.**
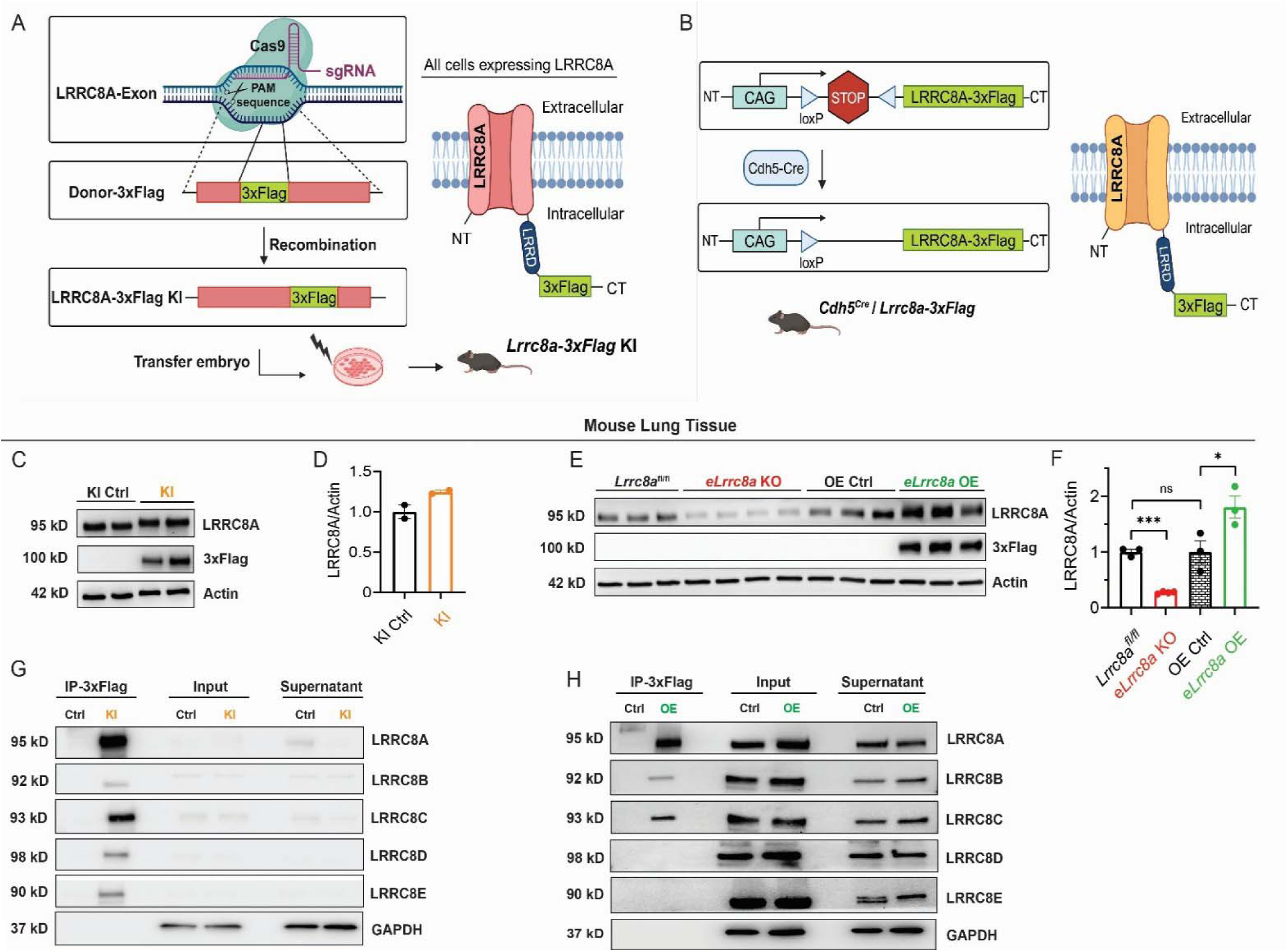
Endothelial LRRC8 complex is a LRRC8A/B/C heteromer. **(A)** Generation of *Lrrc8a*-3xFlag knock-in (KI) mice where a 3xFlag epitope was knocked into the endogenous *Lrrc8a* C-terminal locus. **(B)** Generation of endothelium-targeted *Lrrc8a*-3xFlag overexpression (*eLrrc8a* OE) mice by crossing CAG-LSL-*Lrrc8a*-3xFlag transgenic mice with *Cdh5*-Cre mice to induce endothelial-targeted *Lrrc8a*-3xFlag overexpression upon Cre-mediated excision of floxed STOP codon. **(C, D)** Validation of *Lrrc8a*-3xFlag KI mice by western blot (WB) for LRRC8A and Flag protein expression in lung tissue lysates isolated from *Lrrc8a*-3xFlag KI mice, as compared to wildtype control (KI Ctrl, **C**), with densitometric quantification of LRRC8A/β-actin (**D,** n = 2). **(E, F)** Validation of *eLrrc8a*-3xFlag OE mice by WB for LRRC8A and Flag protein expression in lung tissue lysates isolated from *Lrrc8a^fl/fl^*, *eLrrc8a* KO, CAG-LSL-*Lrrc8a*-3xFlag transgenic (OE Ctrl) and CDH5-Cre;CAG-LSL-LRRC8A-3xFlag mice (*eLrrc8a* OE, **E**), and densitometric quantification of LRRC8A/β-actin (**F,** n = 3). **(G)** Immunoprecipitation (IP) with anti-Flag antibody followed by WB for LRRC8A-E in lung tissue lysates isolated from *Lrrc8a*-3xFlag KI (KI) and control mice (Ctrl). **(H)** IP-WB using lung tissue lysates from *eLrrc8a* OE and control (Ctrl) mice. GAPDH was blotted for each LRRC8 subunit respectively and one representative blot was shown. Statistical significance between the indicated groups in all blots was calculated using a two-tailed unpaired Student’s t-test. Data are represented as mean ± S.E.M. ns - not significant; \**p* < 0.05; \*\*\**p* < 0.001.

### Endothelium exhibit co-dependent LRRC8A/B/C protein levels

We next examined LRRC8A, B, C and D protein levels in endothelium-enriched lung tissue from e*Lrrc8a* KO, *Lrrc8b* KO and *Lrrc8c* KO mice to assess for co-dependent protein levels that might be consistent with the presence of an endothelial LRRC8A/B/C hetero-hexamer suggested by the co-IP data. Interestingly, reductions in LRRC8A protein in e*Lrrc8a* KO lung tissues are associated with significant reductions in LRRC8B and LRRC8C expression but not LRRC8D **(Fig. 4A-E).** Similarly, LRRC8C depletion in *Lrrc8c* knockout lung tissue is associated with marked reductions in LRRC8A and LRRC8B protein levels, with no change in LRRC8D **(Fig. 4F-J).** Curiously, elimination of LRRC8B protein in *Lrrc8b* knockout lung tissues augmented LRRC8A protein, without significant changes in LRRC8C, nor LRRC8D protein levels **(Fig 4K-O).** Indeed, a similar pattern of co-dependent LRRC8 protein levels is observed in HUVECs upon siRNA-mediated LRRC8A/B/C knock-down (KD) (**Fig. S2**). Overall, these data in both primary endothelium and HUVEC demonstrate a co-dependent relationship among LRRC8A, LRRC8B and LRRC8C, but not LRRC8D protein levels, supporting the notion of an endothelial LRRC8A/B/C heteromeric channel complex.

**Figure 4.**
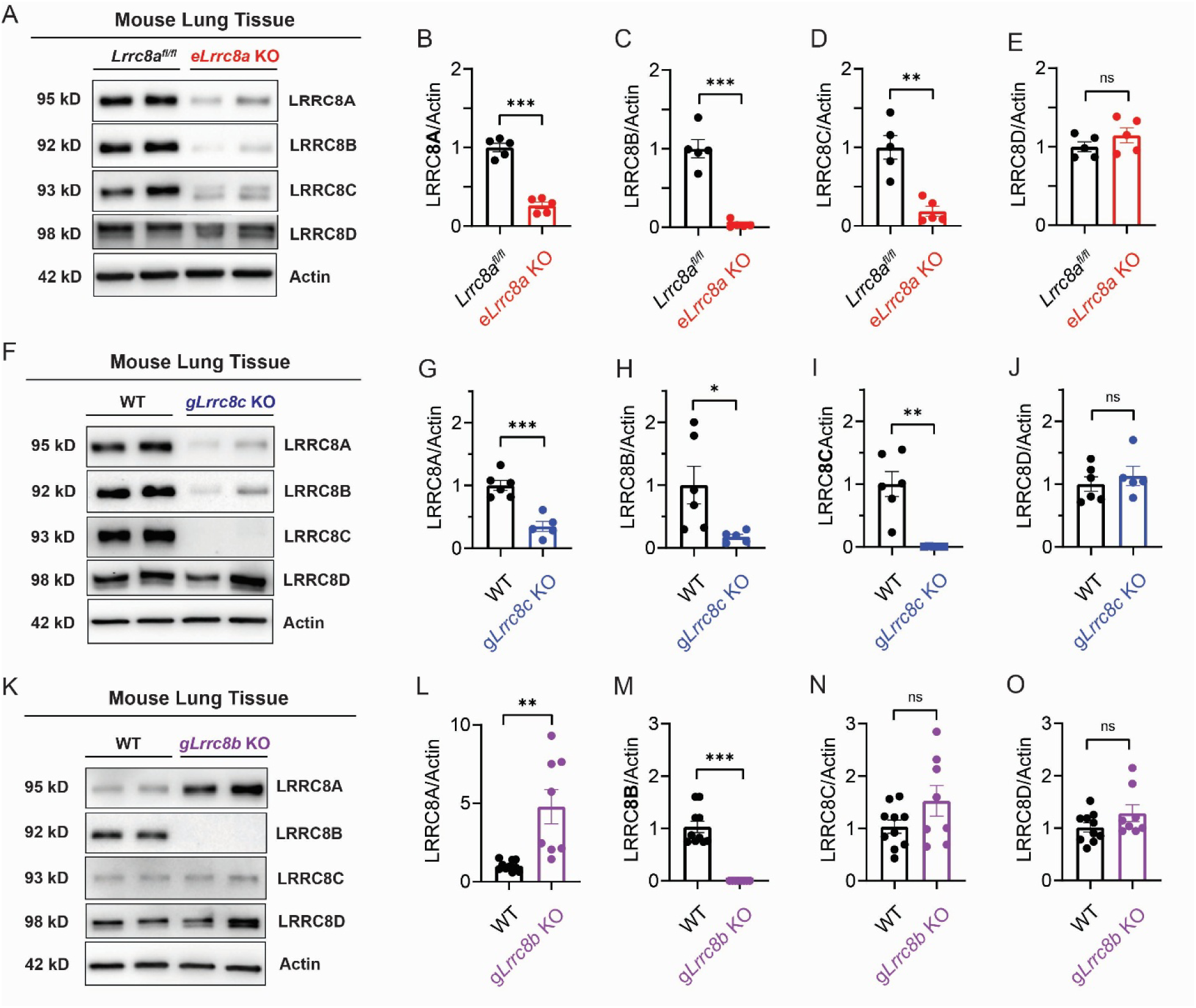
LRRC8A, LRRC8B, and LRRC8C subunits exhibit co-dependent expression in murine endothelium. (A-E) Western blot and quantification of LRRC8A, 8B, 8C and 8D relative to β-actin in lung tissue lysates isolated from *Lrrc8a^fl/fl^*control (n = 5) and *eLrrc8a* KO (n = 5) mice. **(F-J)** Western blot and densitometric quantification of LRRC8A, 8B, 8C and 8D relative to β-actin in lung tissue lysates isolated from wild-type (WT, n = 6) and global *Lrrc8c* knockout (*gLrrc8c* KO, n = 5) mice. **(K-O)** Western blot and denistometric quantification of LRRC8A, 8B, 8C and 8D relative to β-actin in lung tissue lysates from WT (n = 10) and global *Lrrc8b* knockout (*gLrrc8b* KO, n = 8) mice. β-actin was blotted for each LRRC8 subunit respectively (see supplemental materials) and one representative blot was shown. Statistical significance between indicated groups was calculated using a two-tailed unpaired Student’s t-test. Data are represented as mean ± S.E.M. ns - not significant; \**p* < 0.05; \*\**p* < 0.01; \*\*\**p* < 0.001.

### LRRC8C contributes to endothelial VRAC current and Akt/eNOS signaling

Based on the human genetic data implicating LRRC8C as a possible regulator of systolic and diastolic blood pressure (**Fig. 1**), in addition to the biochemical data supporting an LRRC8A/B/C heteromer (**Fig. 3-4, Fig. S1-2**), we next sought to examine the contribution of LRRC8C to both endothelial VRAC (eVRAC) and LRRC8-dependent AKT-eNOS signaling, as we showed previously that both eVRAC and endothelial AKT-eNOS signaling were LRRC8A-dependent ^3^. shRNA-mediated LRRC8C KD resulted in 70% reduction in eVRAC as compared to 85% eVRAC reduction upon LRRC8A KD (**Fig. 5**), indicating that LRRC8C also contributes to LRRC8 channel activity in the endothelium.

**Figure 5.**
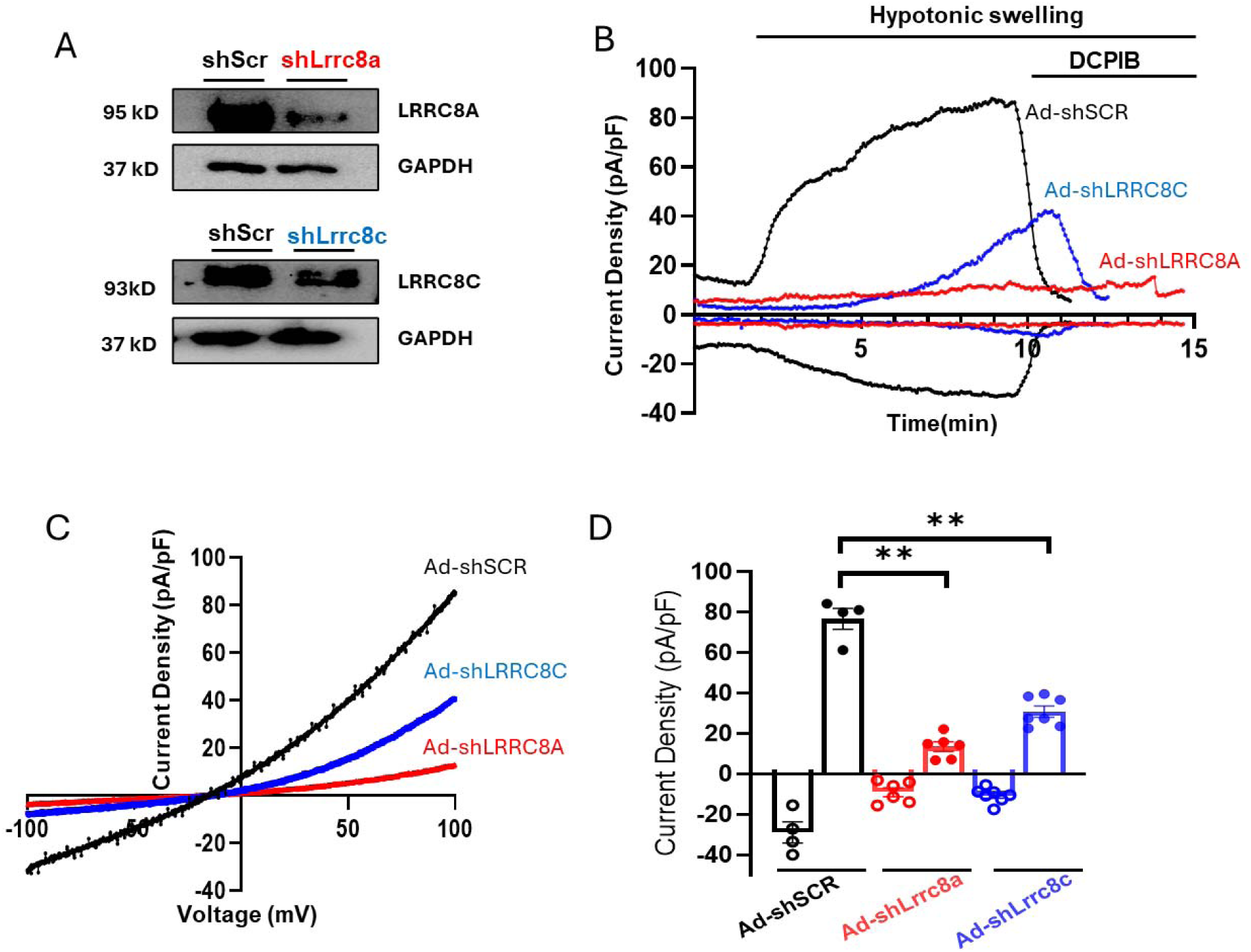
LRRC8A and LRRC8C contribute to Volume-Regulated Anion Channel (VRAC) currents in human umbilical vein endothelial cells (HUVECs). **(A)** Western blot of LRRC8A and LRRC8C in HUVECs transduced with an adenovirus expressing a short-hairpin RNA targetting *LRRC8A* (Ad-sh*Lrrc8a*) or *LRRC8C* (Ad-sh*Lrrc8c*), as compared to cells transduced with scrambled control short-hairpin RNA (Ad-shScr). **(B)** Representative current-time relationship of hypotonically (210 mOsm) induced VRAC currents in HUVECs transduced with Ad-shScr, Ad-sh*Lrrc8a* or Ad-sh*Lrrc8c*, followed by application of 10 µM of the VRAC inhibitor DCPIB. **(C)** Current-voltage relationship of VRAC currents during voltage ramps from −100 mV to +100 mV after hypotonic swelling of HUVECs transduced with Ad-shScr, Ad-sh*Lrrc8a* or Ad-sh*Lrrc8c*. **(D)** Mean outward (at +100 mV) and inward (at −100 mV) current densities in HUVECs transduced with Ad-shScr (n = 4), Ad-sh*Lrrc8a* (n = 6) or Ad-sh*Lrrc8c* (n = 7). Statistical significance between indicated groups was calculated using a two-tailed, unpaired Student’s t-test. Data are represented as mean ± S.E.M. \*\**p* < 0.01.

We next examined AKT-eNOS signaling in primary endothelium using lung tissue from WT, e*Lrrc8a* KO, *Lrrc8c* KO, and *Lrrc8b* KO mice. Basal levels of AKT-eNOS signaling in e*Lrrc8a* KO lungs recapitulates our previous findings in human umbilical vein endothelial cells (HUVECs) with siRNA induced LRRC8a knockdown ^3^, as evidenced by reductions in p-eNOS, p-AKT1 and p-AKT2 phosphorylation (**Fig. 6A-H**). These data also validate the use of lung tissue for evaluation of basal AKT-eNOS signaling from primary endothelium. Similarly, LRRC8C ablation in lung tissues (**Fig. 6I-P**) and HUVECs (**Fig. S3A-G**) replicates the reductions in pAKT1, pAKT2, and p-eNOS signaling observed with LRRC8A depletion in the endothelium (**Fig. 6A-H**) ^3^.

**Figure 6.**
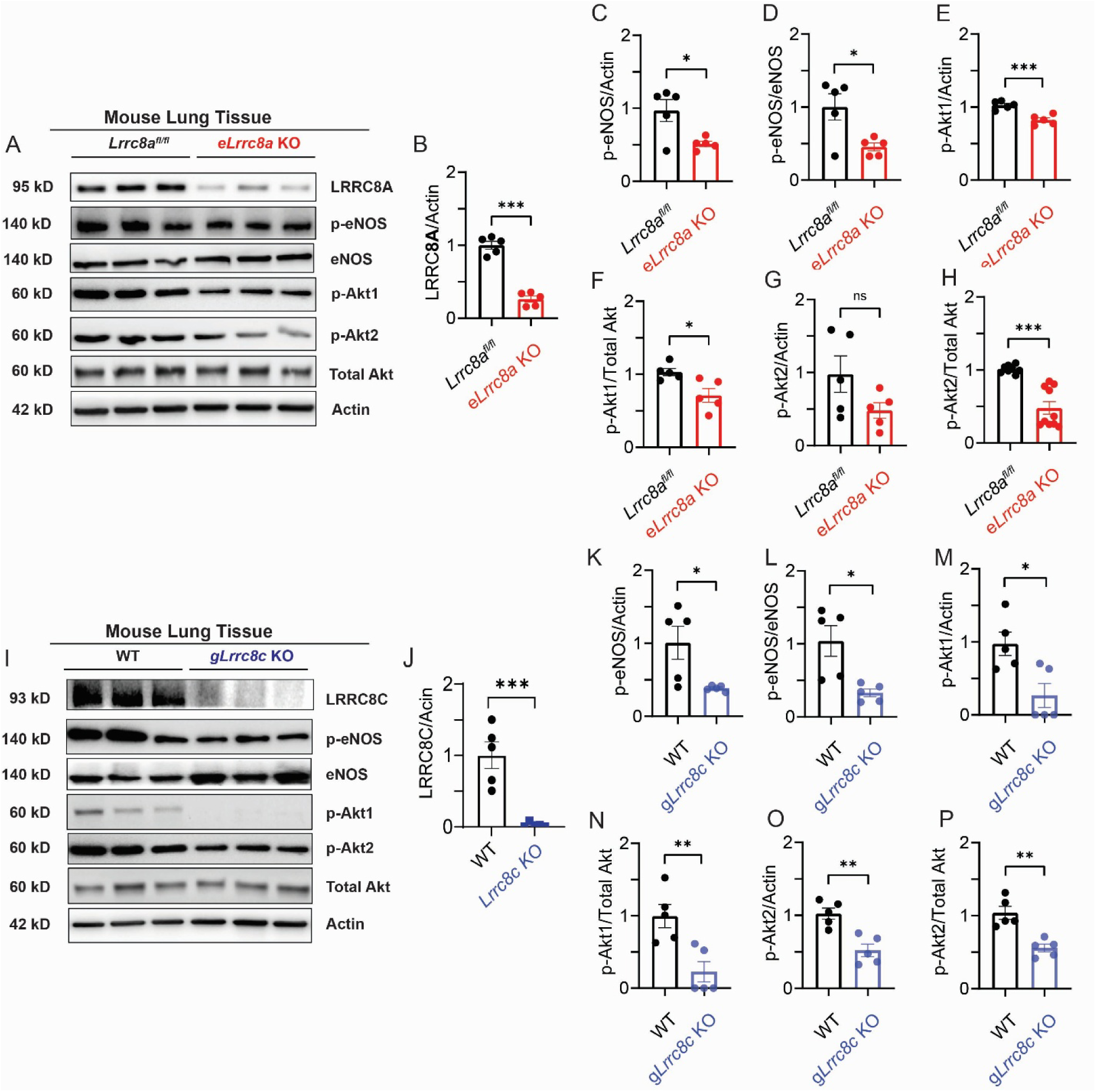
LRRC8C regulates endothelial Akt-eNOS signaling. **(A)** Western blots of LRRC8A, phosphorylated eNOS (p-eNOS), total eNOS, p-Akt1, p-Akt2 and total Akt in *Lrrc8a^fl/fl^* and e*Lrrc8a* KO mice. **(B-H)** Quantification of LRRC8A/β-actin, p-eNOS/total eNOS, p-eNOS/β-actin, pAkt1/total Akt1, pAkt1/β-actin, pAkt2/total Akt2 and pAkt2/β-actin (n = 5). (**I**) Western blots of p-eNOS, total eNOS, pAkt1, pAkt2 and total Akt in wildtype (WT) and *gLrrc8c* KO mice. **(J-P)** Quantification of LRRC8C/β-actin, p-eNOS/total eNOS, p-eNOS/β-actin, pAkt1/total Akt1, pAkt1/β-actin, pAkt2/total Akt2, and pAkt2/β-actin (n = 5). Statistical significance between indicated groups was calculated using a two-tailed unpaired Student’s t-test. Data are represented as mean ± S.E.M. \**p* < 0.05; \*\**p* < 0.01; \*\*\**p* < 0.001.

### e*Lrrc8a* KO and *Lrrc8c* KO mice exhibit increased myogenic tone and impaired endothelial nitric oxide dependent vasodilation

To determine if impaired endothelial AKT-eNOS signaling observed upon LRRC8A and LRRC8C ablation also abrogated vascular function, we used third order mesenteric arteries mounted in a pressure myograph to measure myogenic tone and vascular relaxation. The vessel diameter was monitored after stepwise increases in intraluminal pressure. Mesenteric arteries of WT mice exhibited normal myogenic tone where the vessel diameter increased proportionally with pressure until ∼60 mmHg, and active myogenic constriction was detected with maintenance of a constant pressure– diameter relationship between 60 and 120 mmHg. In contrast to WT mice, resistance arteries from both e*Lrrc8a* KO mice and g*Lrrc8c* KO mice exhibited significantly increased myogenic response, initiating active constrictions from 40 mmHg of pressure **(Fig. 7A, B, G, H).** Importantly, the differences in active diameter changes were abolished when endothelium was denuded from the third order mesenteric artery preparations **(Fig. 7C, D)**, consistent with an endothelium-dependent mechanism underlying modulation of myogenic tone by LRRC8A and LRRC8C. All vessels had similar maximum calcium-independent passive diameters, with no statistically significant difference between WT, e*Lrrc8a* KO and g*Lrrc8c* KO **(Fig. 7E, F)**. In addition, pressure myograph recordings performed on mesenteric arteries from endothelial specific *Lrrc8a* overexpression (e*Lrrc8a* OE) mice revealed unchanged active and passive diameter as well as myogenic tone compared to their 3xFlag wildtype controls **(Fig. S4A-C)**.

**Figure 7.**
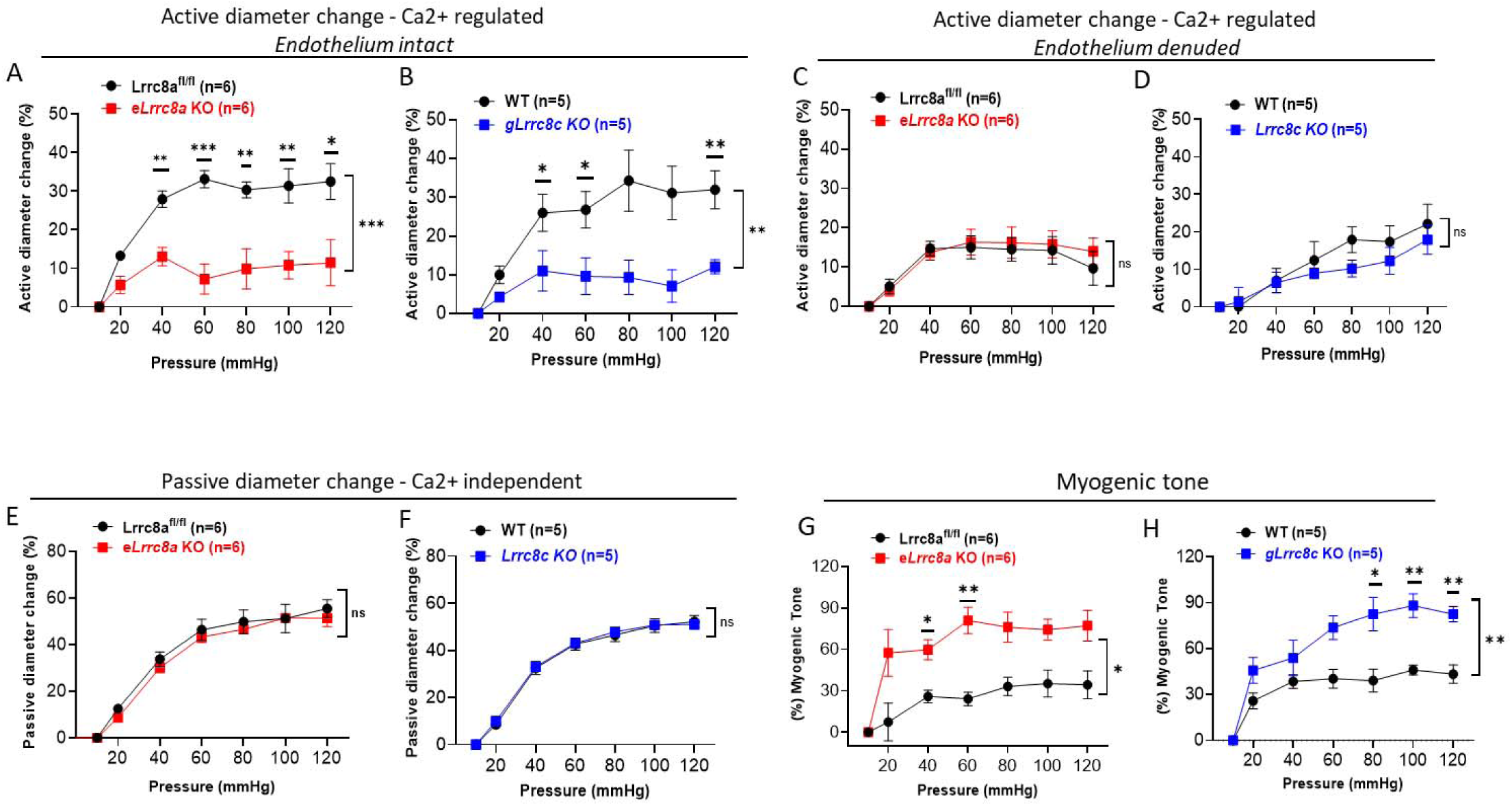
Endothelial dependent increase in myogenic tone of endothelial-specific *Lrrc8a* knockout and global *Lrrc8c* knockout mesenteric arteries as measured by pressure myography. (**A, B**) Active diameter changes of cannulated third order mesenteric arteries with intact endothelium in calcium-present conditions during stepwise increments of pressure comparing eLLRC8a KO and g*Lrrc8c* KO mice to their respective controls. **(C, D)** Active diameter changes of cannulated third order mesenteric arteries with denuded endothelium during stepwise increments of pressure comparing e*Lrrc8a* KO and g*Lrrc8c* KO mice to their respective controls. (**E, F)** Passive diameter changes in mesenteric arteries isolated from *eLrrc8a* KO and *gLrrc8c* KO mice in calcium-free conditions, as compared to their respective controls. **(G, H)** Myogenic tone calculated as the relative difference in active and passive diameters in endothelium-intact mesenteric arteries. n=6 in *Lrrc8a^fl/fl^* and e*Lrrc8a* KO. n=5 in WT and g*Lrrc8c* KO. Statistical significance was determined by two-way ANOVA with Bonferroni post-hoc test. Data are reresented as mean ± S.E.M. ns - not significant; \**p* < 0.05; \*\**p* < 0.01; \*\*\**p* < 0.001.

These results are consistent with prior observations that LRRC8A overexpression does not increase VRAC currents above WT levels, and may result in suppression of endogenous VRAC currents ^5,7^. Similar results were obtained from *Lrrc8b* KO mice **(Fig. S4 D-F)**, with only a mild, but statistically significant reduction in observed passive diameter change.

As both LRRC8A and LRRC8C regulate AKT-eNOS signaling in endothelium and nitric oxide is critical for endothelial-dependent vasodilation ^13^, we next examined the impact of LRRC8A and LRRC8C ablation on nitric oxide-mediated vasodilation in mesenteric arteries. e*Lrrc8a* KO and *Lrrc8c* KO mesenteric arteries responded similarly to 10 μM phenylephrine pre-constriction, but exhibited impaired relaxation in response to the endothelium-dependent vasodilator, acetylcholine (ACh, 1 nM-1 mM) compared to their respective wildtype controls (**Fig. 8A, B)**. In contrast, endothelium-independent vasorelaxation upon papaverine (1 nM-1 mM) treatment, which induces cAMP and cGMP production via inhibition of phosphodiesterase^14^, is not affected by the absence of LRRC8A or LRRC8C **(Fig. 8C, D)**, suggesting impaired LRRC8A/C signaling in endothelium specifically. Moreover, eNOS inhibition via addition of L-N^G^-Nitro arginine methyl ester (L-NAME, 10μM) led to a significant reduction in vasodilation in response to acetylcholine in wildtype **(Fig. S5A, C)** but not e*Lrrc8a* KO **(Fig. S5B)** or *Lrrc8c* KO mesenteric arteries **(Fig. S5D),** indicating decreased NO bioavailability in e*Lrrc8a* KO and *Lrrc8c* KO vessels. In contrast, *gLrrc8b* KO mesenteric arteries failed to show changes in vasodilatory responses to acetylcholine or papaverine **(Fig. S5E, F)**.

**Figure 8.**
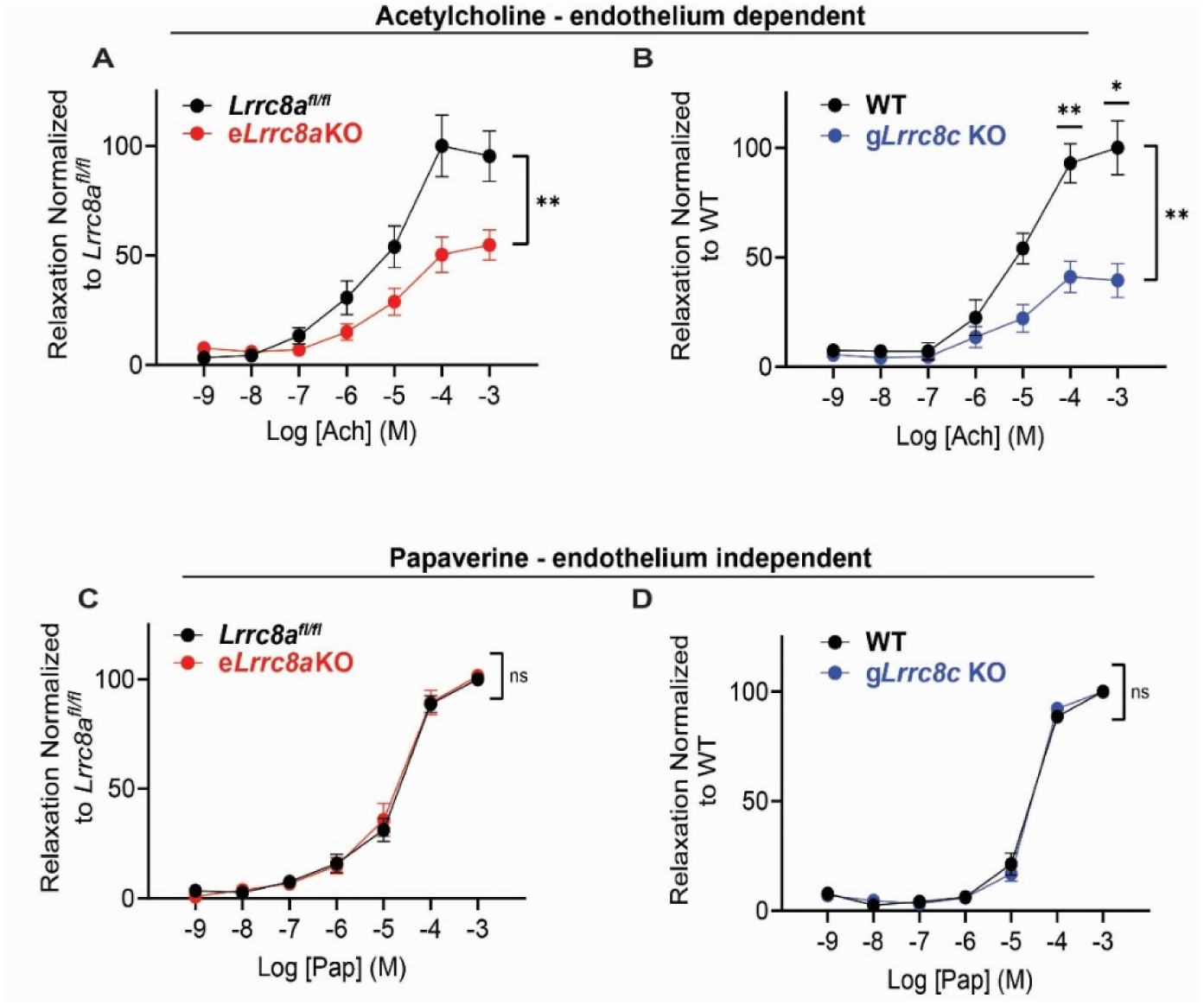
Impaired acetylcholine-mediated vasodilation in mesenteric arteries of endothelial-specific *Lrrc8a* knockout and global *Lrrc8c* knockout mice. (**A, B**) Dose-response curves showing relaxation of third order mesenteric arteries in response to increasing doses of acetylcholine (Ach) in e*Lrrc8a* KO and g*Lrrc8c* KO compared to their respective controls. **(C, D)** Relaxation of mesenteric arteries in response to increasing doses of papaverine (Pap) in *eLrrc8a* KO and and *gLrrc8*c KO compared to their respective controls. n=7 in *Lrrc8a^fl/fl^*and e*Lrrc8a* KO. n=6 in WT and g*Lrrc8c* KO. Statistical significance was determined by two-way ANOVA with Bonferroni post-hoc test. Data are represented as mean ± S.E.M. ns - not significant; \**p* < 0.05; \*\**p* < 0.01; \*\*\**p* < 0.001.

### Loss of LRRC8C exacerbates angiotensin II-induced hypertension

To determine how the observed changes in myogenic tone and endothelial-dependent vasodilatory responses in resistance vessels affects systemic blood pressure, we next measured tail cuff blood pressures in *Lrrc8c* and *Lrrc8b* KO mice. Neither *Lrrc8b* nor *Lrrc8c* KO mice exhibited changes in systolic or mean blood pressures compared to their respective wild-type control under basal conditions **(Fig S6A,C,D,F)**, similar to our previous observations in e*Lrrc8a* KO mice ^3^. However, *Lrrc8c* KO mice had mildly elevated diastolic blood pressure **(Fig. S6B,** *p*=0.0499). To test for a propensity for developing hypertension, we implanted angiotensin II infusing osmotic minipumps into *Lrrc8c* KO and WT control mice. After 4 weeks of mild angiotensin II infusion, *Lrrc8c* KO mice developed exacerbated hypertension as compared to Ang II-treated WT mice (**Fig 9A-C**). There was no gender difference in blood pressure response to Angiotensin II chronic infusion in *Lrrc8c* KO and their wildtype control mice (**Fig S7A-D**). These data are consistent with the association between *LRRC8C* and hypertension in human GWAS and PheWAS studies, and highlights the shared function of LRRC8C and LRRC8A in regulating endothelial LRRC8 complex expression, VRAC current, as well as coordinating endothelial Akt/eNOS signaling to impact myogenic tone and vasodilatory activity in resistance arteries and regulation of systemic blood pressure.

**Figure 9.**
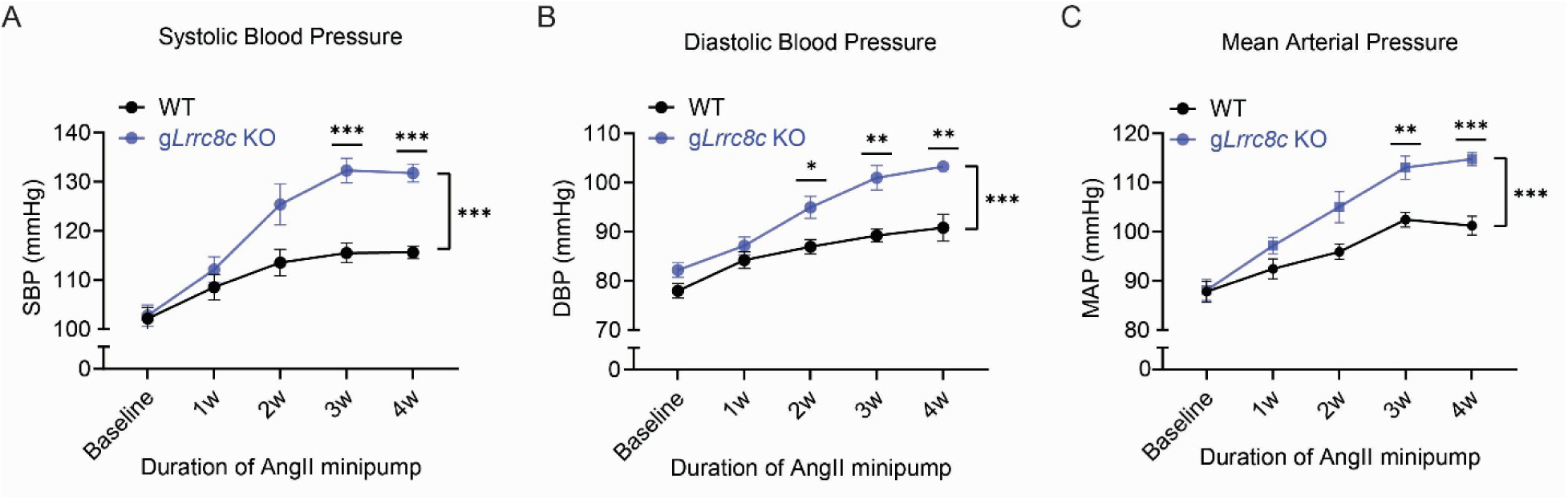
Loss of LRRC8C exacerbates angiotensin II-induced hypertension. Tail-cuff systolic blood pressures **(A)**, diastolic blood pressure **(B)**, and mean arterial pressure **(C)** of wildtype (WT, n = 10 males and 7 females) and g*Lrrc8c* KO (n = 9 males and 7 females) mice after 4 weeks of chronic angiotensin-II infusion. Statistical significance was determined by two-way ANOVA with Bonferroni post-hoc test. Data are represented as mean ± S.E.M. * *p* < 0.05; \*\**p* < 0.01; \*\*\**p* < 0.001.

## Discussion

Here we show the LRRC8 complex in the endothelium is largely comprised of LRRC8A/B/C heteromers. LRRC8C is the second dominant functional subunit within the endothelial LRRC8A/B/C complex. LRRC8C depletion phenocopies endothelial LRRC8A depletion by reducing VRAC currents, compromising AKT-eNOS signaling, increasing myogenic tone, and impairing eNOS dependent vasodilation, which ultimately exacerbates angiotensin-induced hypertension. These findings suggest a non-redundant role for LRRC8C in regulating endothelial AKT/eNOS pathway, vascular relaxation and susceptibility to hypertension.

The identification of SNP rs12032393 on nucleotide 1:89713973 (GRCh38) in association with a reduction in diastolic blood pressure is interesting and clinically relevant^15^, given that the African American population demonstrates a remarkably lower frequency of this missense SNP risk allele but a higher prevalence of hypertension^16^. This is consistent with our finding in mice with LRRC8C depletion which worsened hypertension after chronic Angiotensin II infusion. Of note, this SNP confers a N468S mutation in *LRRC8C*, which translates to a less polar hydroxyl side chain in serine replacing a polar amide side chain in asparagine in leucine-rich repeat domains. This could be important in protein-protein binding by establishing electrostatic bonds with the docking protein, or may indirectly affect channel activation.

Indeed, we have previously demonstrated a constitutive LRRC8A interaction with adaptor protein GRB2 (growth factor receptor-bound 2) and form a LRRC8A-GRB2-Cav1 signaling complex which regulates PI3K/AKT signaling in adipocytes, skeletal myotubes and endothelium ^3,10,17,18^. Given the functional similarity with LRRC8A, LRRC8C may share the same protein binding structure to regulate endothelial Akt-eNOS signaling. Therefore, further investigation of the in vivo functional impact of the N468S mutation in *LRRC8C on vascular outcome and its underlying mechanism* is warranted, though beyond the scope of current study.

It is also noteworthy that endothelial cells are known to release adenosine triphosphate (ATP) in response to shear stress, which serves as an important extracellular ligand of autocrine and paracrine signaling^19^. In addition to classical exocytosis, non-vesicular endothelial ATP release mechanisms have been demonstrated via ATP-permeable pannexin1 channels ^20^ which shares anion permeability and a large ion-conducting pore, and is indirectly mechanically-activated via Piezo1-mediated Ca^2+^ influx ^21^. This ATP release activates purinergic P2Y2 receptors in endothelial cells, induces downstream Akt/eNOS signaling, and stimulates nitric oxide (NO) which reduces vascular tone and blood pressure ^19,21^. Therefore, it is also possible that LRRC8 regulation of AKT-eNOS signaling is mediated by LRRC8 channels functioning as endothelial ATP release channels, as reported by others in microglia and macrophages ^22,23^. It is notable that LRRC8C knockdown in HUVECs reduces the inward VRAC component proportionately more than the outward component, as compared to LRRC8A KD, resulting in a change in rectification ratio. This preferential reduction in inward VRAC current in LRRC8C KD HUVECs may reflect a change in channel permeability, possibly ATP permeation.

LRRC8C was previously reported to function as a non-essential component of the VRAC, and plays a redundant role in the efflux of amino acids, such as aspartate and glutamate, in response to osmotic stress^24^. Recent studies have pointed to a non-redundant role of LRRC8C in mediating multiple pathophysiological processes. In heteromeric complexes with LRRC8A and LRRC8E, LRRC8C acts as a broadly expressed transporter of 2’3’-cyclic-GMP-AMP (cGAMP) ^25^, which is an immunotransmitter released from diseased cells and taken up by host cells to initiate the stimulator of interferon genes (STING) immune response ^26^. Concepcion et.al, further demonstrated that LRRC8C deletion abolishes VRAC currents and exacerbates T cell-dependent immune responses via cGAMP-STING signaling ^27^, suggesting that LRRC8C is a critically functional component in the LRRC8A/C/E hetero-hexamers in T-cells, similar to our findings of the role of LRRC8C in endothelial LRRC8A/B/C heteromer in endothelial cells. Also of note, two de novo variants of *LRRC8C* were recently reported in two unrelated human individuals who presented with an identical but novel syndromic multisystem disorder ^28^. Both *LRRC8C* mutations resulted in constitutive activation of VRAC in isotonic conditions when wildtype channels are closed. The consequent functional disturbance is associated with prominent vascular involvement in both patients including marked skin telangiectasia and gastrointestinal vascular dysplasia, which highlighted the role of LRRC8C in regulating endothelial function. These findings support our *Lrrc8c* loss-of-function studies in endothelial cells where increased myogenic tone and impaired vasodilatation were observed.

It is intriguing that endothelial specific overexpression of LRRC8A did not alter vascular reactivity as opposed to an expected gain-of-function effect. Interestingly, LRRC8A overexpression has been previously shown to cause an unexpected suppression of endogenous VRAC currents ^5,7^. Although LRRC8A and LRRC8C co-expression did not suppress *VRAC current, n*either LRRC8A and LRRC8C co-expression, nor any other combination tested, significantly increased current amplitudes above wildtype values ^5^. These findings support the notion that a specific stoichiometry of LRRC8 subunits is required to form functional VRAC in different organ and cell types, and different combinations of LRRC8B-E plus LRRC8A yield VRAC currents with different inactivation kinetics, rectification and single-channel conductance ^7^.

Our study is limited by the availability of endothelial specific LRRC8C knockout mice in the interpretation of the in vivo hypertensive response to Angiotensin II. However, the remarkable changes in myogenic tone of endothelium intact vs denuded mesenteric arteries strongly supports a critical role of LRRC8C in endothelium specific signaling. This is also corroborated by the vasodilatory study using endothelium-independent vasodilator as well as an eNOS inhibitor. Because LRRC8C is selectively enriched in endothelial cells and its deletion is not lethal in mice, in contrast to the deletion of the obligatory VRAC channel subunit LRRC8A, pharmacological targeting of LRRC8C may represent a specific and safer approach to various vascular diseases which involves endothelial dysfunction.

## Acknowledgments

We gratefully acknowledge Dr. Thomas Jentsch (Leibniz-Forschungsinstitut für Molekulare Pharmakologie, Berlin, Germany) and Dr. Axel Concepcion (University of Chicago, Chicago, IL) for kindly providing rabbit polyclonal antibodies against LRRC8B, LRRC8C, LRRC8D, and LRRC8E. We thank the Genome Engineering & Stem Cell Center (GESC@MGI) at the Washington University in St. Louis for reagent validation services to generate the *Lrrc8a*-3xFlag knock-in mice, and g*Lrrc8b* KO mice. Dr. Axel Concepcion for sharing the g*Lrrc8c* KO mice.

## Supplemental Materials

### Supplemental figures and figure legends

**Supplemental Figure 1.**
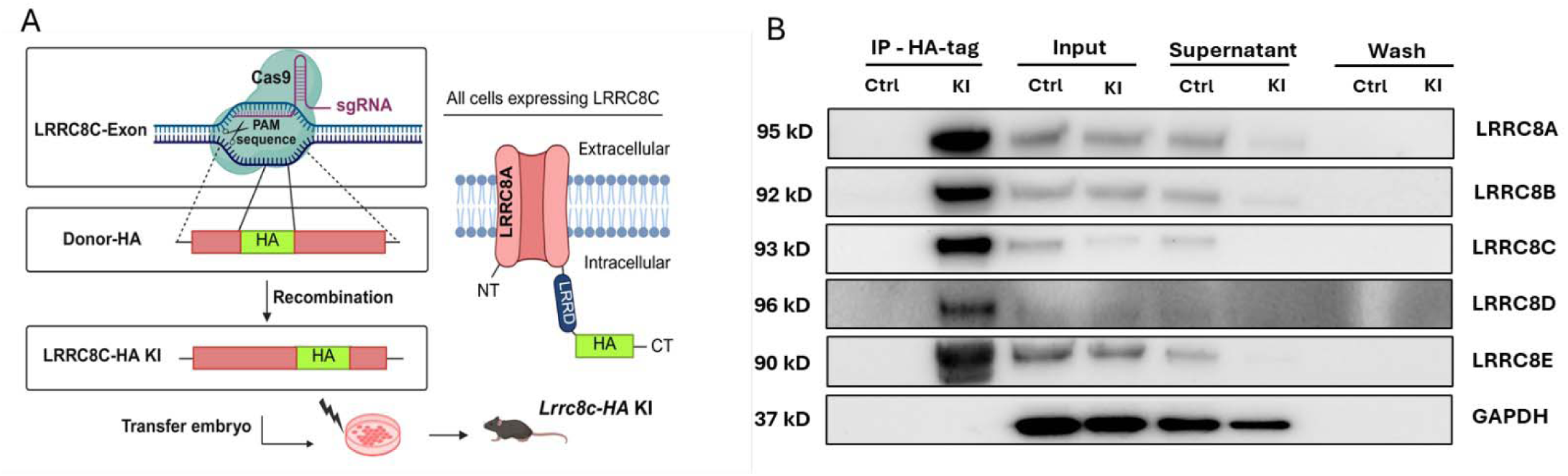
Immunoprecipitation (IP) with anti-HA antibody followed by WB for LRRC8A-E in lung tissue lysates isolated from control mice (Ctrl) and *Lrrc8c*-HA knock-in (KI) mice where a HA epitope was knocked into the endogenous *Lrrc8c* C-terminal locus. GAPDH was blotted for each LRRC8 subunit respectively and one representative blot was shown.

**Supplemental Figure 2.**
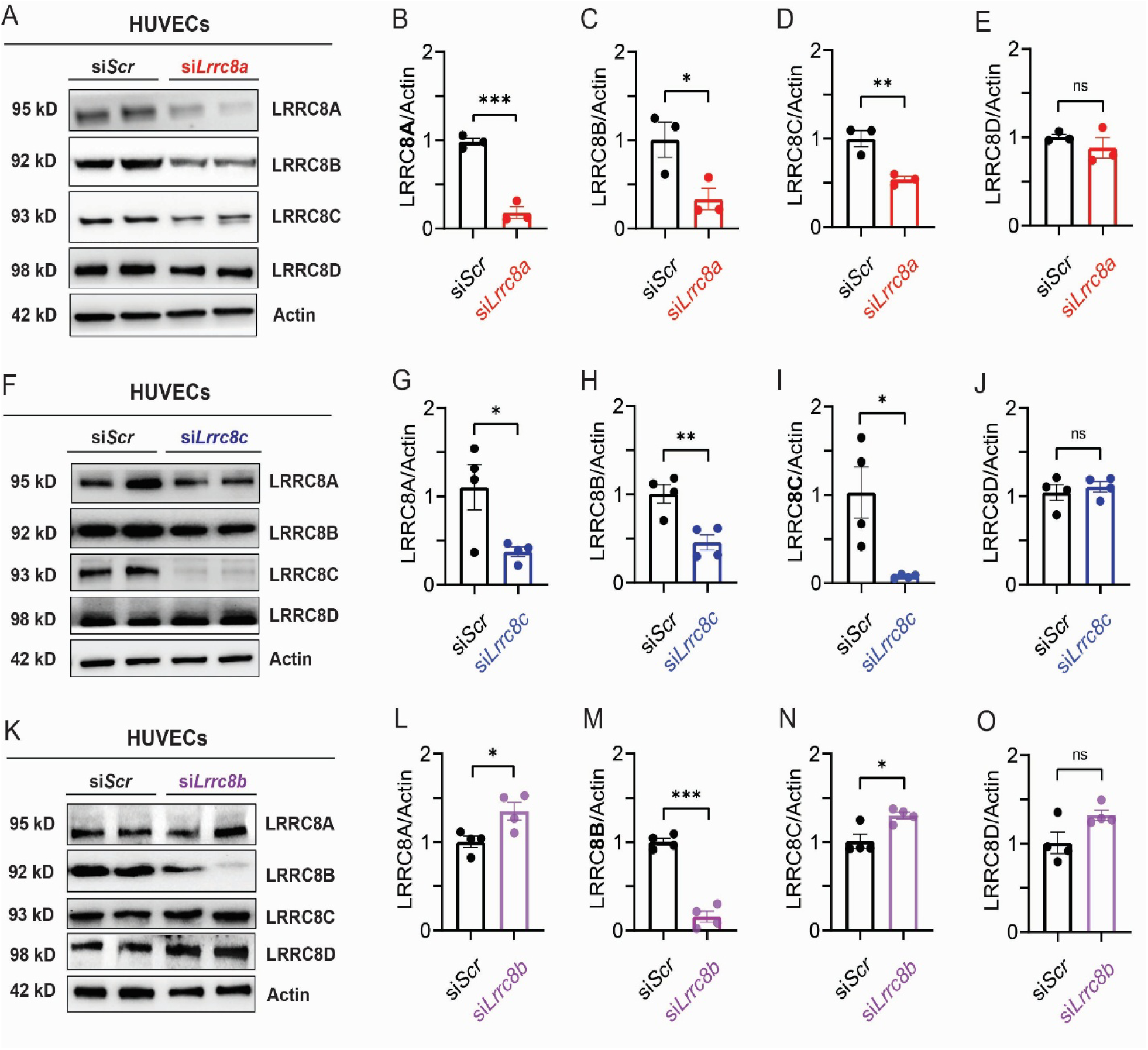
L**R**RC8A**/B/C demonstrate co-dependent expression in human umbilical vein endothelial cells (HUVECs).** (**A-E**) Western blots and densitometric quantification of LRRC8A, 8B, 8C and 8D in scramble control siRNA (siScr) vs *Lrrc8a* siRNA (si*Lrrc8a*) transduced HUVECs (n = 3). (**F-J**) Western blots and quantification of LRRC8A, 8B, 8C and 8D in scramble control siRNA (siScr) vs *Lrrc8c* siRNA (si*Lrrc8c*) transduced HUVECs (n = 4). (**K-O**) Western blots and quantification of LRRC8A, 8B, 8C and 8D in scramble control siRNA (siScr) vs *Lrrc8b* siRNA (si*Lrrc8b*) transduced HUVECs (n = 4). β-actin was blotted for each LRRC8 subunit respectively (see supplemental materials) and one representative blot was shown. n= 3 independent experiments. Statistical significance between indicated groups was calculated using a two-tailed unpaired Student’s t-test. Data are represented as mean ± S.E.M. ns - not significant; *p<0.05; **p<0.01; ***p<0.001.

**Supplemental Figure 3.**
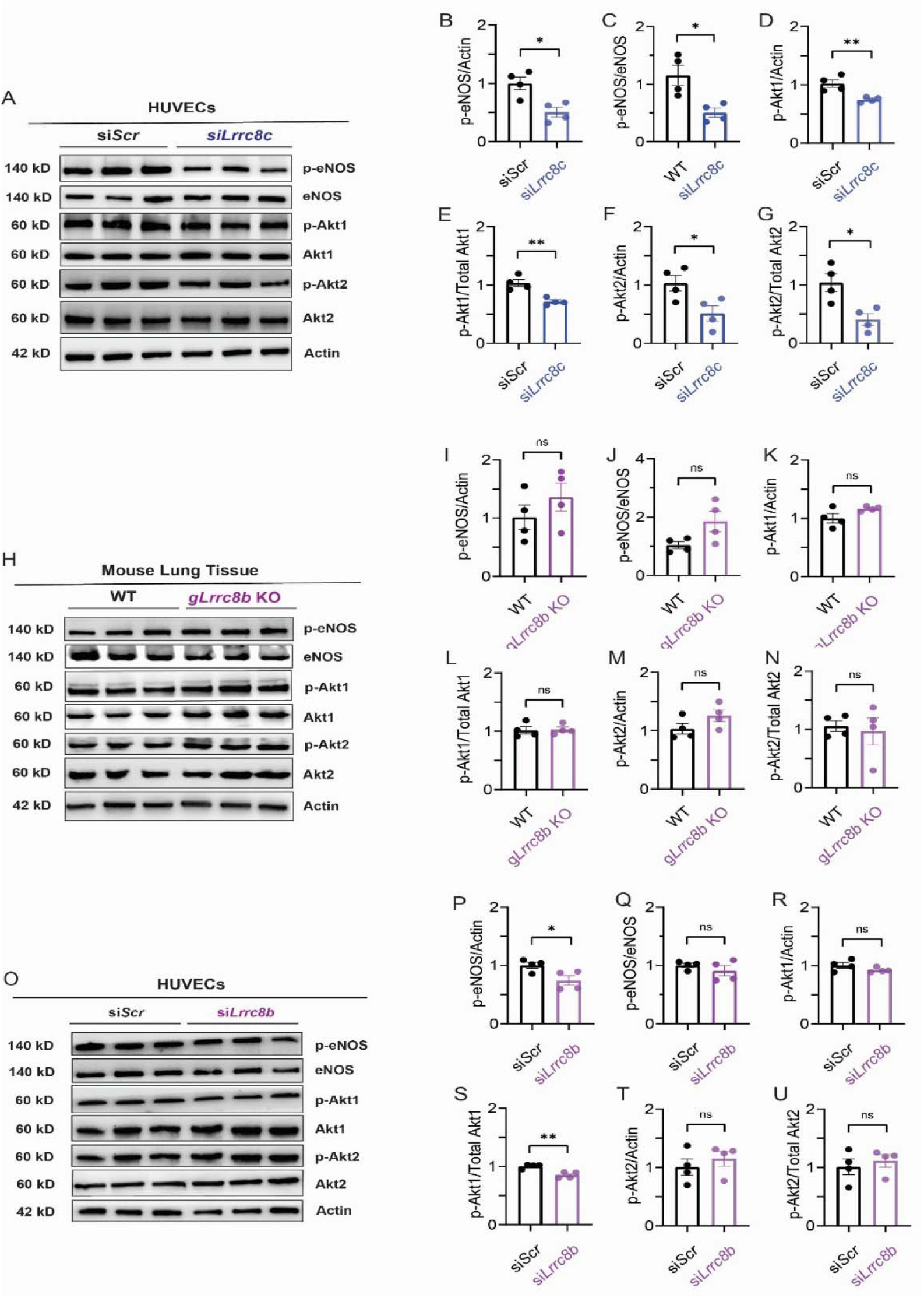
Akt-eNOS signaling is unchanged upon endothelial LRRC8B depletion. **(A)** Western blots of p-eNOS, eNOS, pAkt1, Akt1, pAkt2, Akt2, and β-actin in scramble control siRNA (si-SCR) vs *Lrrc8b* siRNA transduced human umbilical vein endothelial cells (HUVECs) after 30 min of serum starvation. **(B-D)** Quantification of p-eNOS/total eNOS, p-eNOS/β-actin, pAkt1/total Akt1, pAkt1/β-actin, and pAkt2/total Akt2, pAkt2/β-actin. **(E-H)** Western blots with quantification of p-eNOS, eNOS, pAkt1, pAkt2, total Akt, and β-actin in lung tissues from wildtype control (WT) vs global *Lrrc8b* knockout (*Lrrc8b* KO) mice. n = 3 independent experiments. Statistical significance between indicated groups was calculated using a two-tailed unpaired Student’s t-test. Data are represented as mean ± S.E.M. ns - not significant; \**p* < 0.05; \*\**p* < 0.01.

**Supplemental Figure 4.**
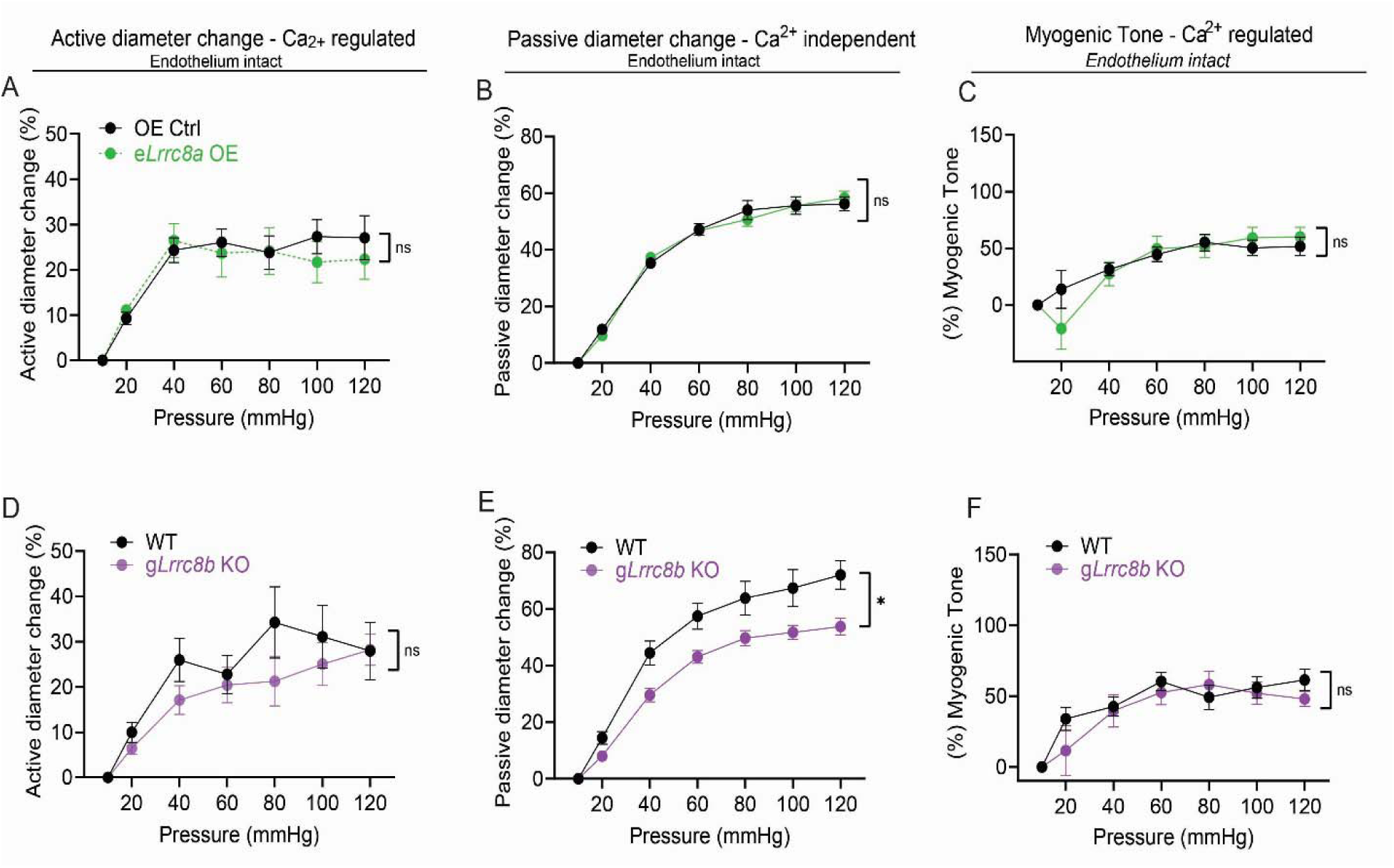
Myogenic tone is unchanged in mesenteric arteries of e*Lrrc8a* KO and g*Lrrc8b* KO. **(A)** Active diameters of cannulated third order mesenteric arteries of e*Lrrc8a* OE mice vs their 3xFLAG control (OE Ctrl) were analyzed during stepwise increments of pressure with intact endothelium. (**B)** Passive diameter was measured in calcium-free conditions with intact endothelium. **(C)** Myogenic tone is assessed by measuring relative difference in active and passive diameters. **(D-F)** Active diameters, passive diameters and myogenic tone were measured in third order mesenteric arteries of g*Lrrc8b* KO mice vs their wildtype control (WT). n=7 in WT and e*Lrrc8a* OE. n=5-6 in WT and g*Lrrc8b* KO. Statistical significance was determined by two-way ANOVA with Bonferroni post-hoc test. Data are represented as mean ± S.E.M. ns - not significant; \**p* < 0.05; \*\**p* < 0.01; \*\*\**p* < 0.001.

**Supplemental Figure 5.**
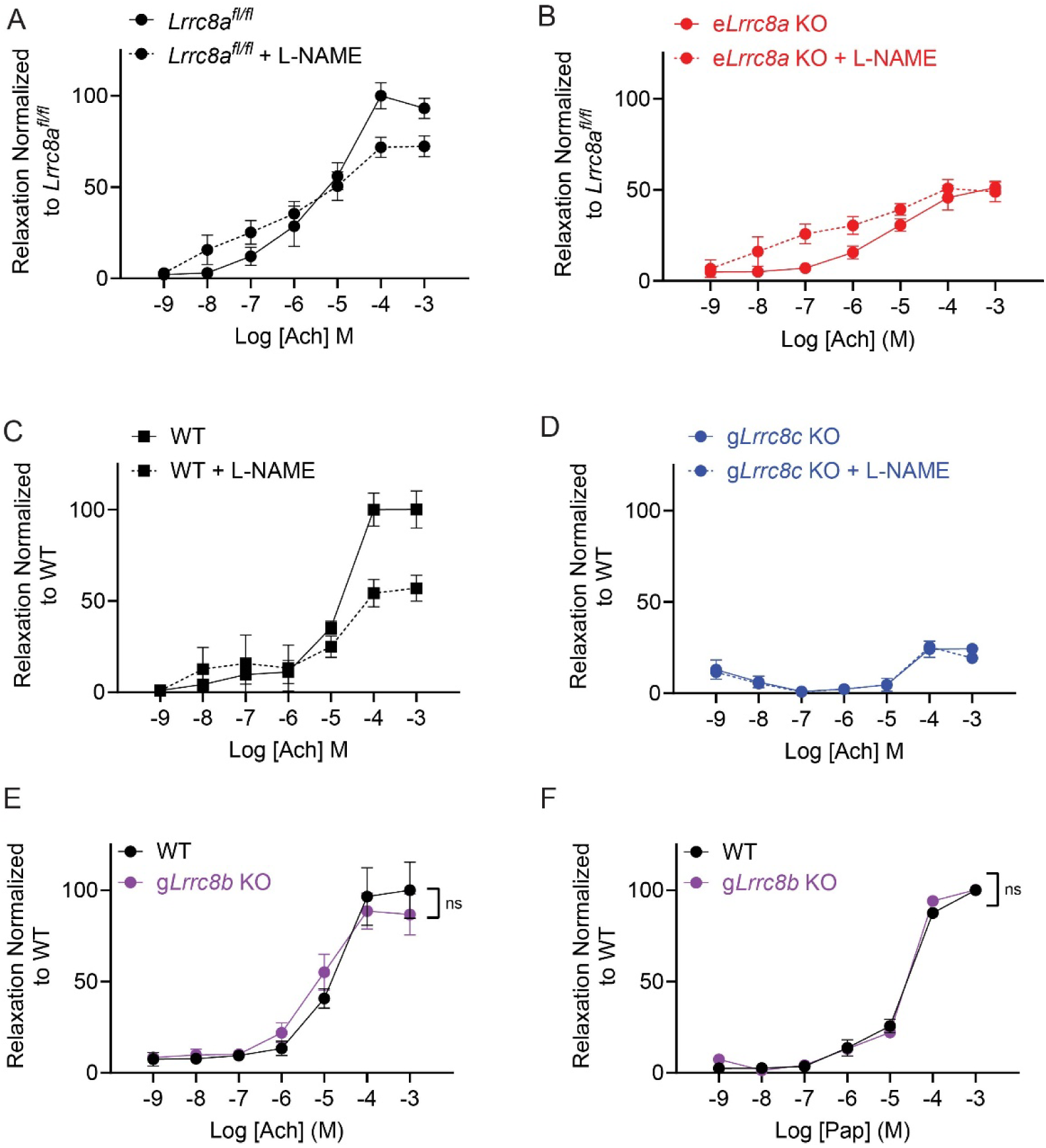
eNOS inhibition mediates impaired relaxation in mesenteric arteries of e*Lrrc8a* KO and g*Lrrc8c* KO. (**A-B**) Relaxation of third order mesenteric arteries from e*Lrrc8a* KO and their *Lrrc8a*^fl/fl^ control mice to increasing doses of acetylcholine (Ach) in the presence of the eNOS inhibitor, L-NAME 10μM. **(C-D)** Relaxation of mesenteric arteries from g*Lrrc8c* KO and their wildtype (WT) control mice to increasing doses of acetylcholine (Ach) in the presence of L-NAME 10μM. (**E-F**) Dose-response curves showing relaxation of third order mesenteric arteries to increasing doses of acetylcholine (Ach) and papaverine (Pap) in g*Lrrc8b* KO mice compared to their WT controls. n=4 in *Lrrc8a*-fl/fl and e*Lrrc8a* KO. n=3 in WT and g*Lrrc8c* KO. n=6 in WT and n=9 in g*Lrrc8b* KO. Statistical significance was determined by two-way ANOVA with Bonferroni post-hoc test. Data are represented as mean ± S.E.M. ns - not significant.

**Supplemental Figure 6.**
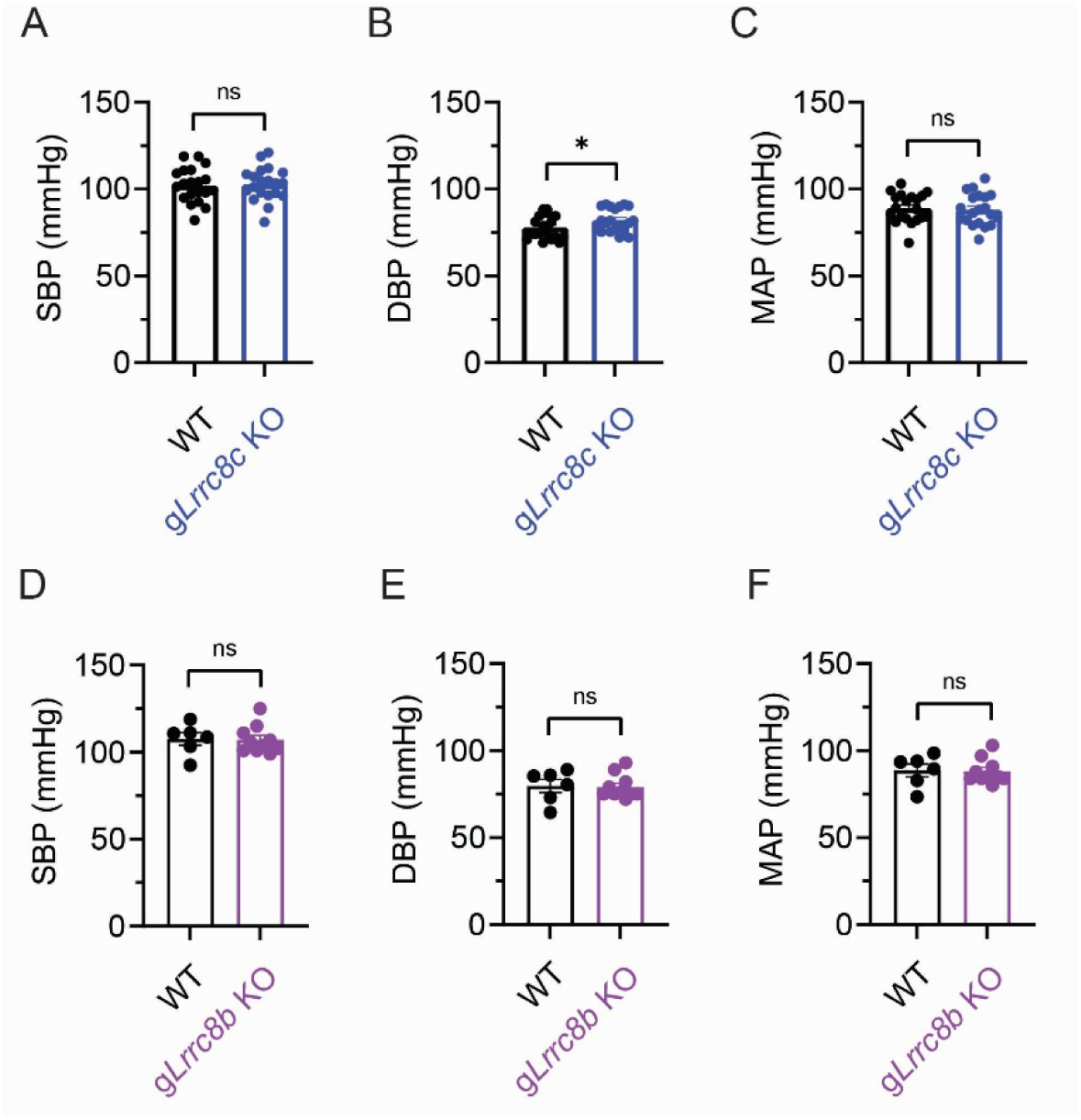
Baseline blood pressures in g*Lrrc8b* KO and g*Lrrc8c* KO mice. (A-C) Tail-cuff systolic blood pressures (SBP), diastolic blood pressure(DBP) and mean arterial pressure (MAP) in *Lrrc8c* KO mice compared to their WT controls at baseline. (**D-F**) SBP, DAP and MAP in global *Lrrc8b* KO and their WT control mice. Statistical significance was determined by two-tailed unpaired Student’s t-test. Data are represented as mean ± S.E.M. ns - not significant; \**p* < 0.05.

**Supplemental Figure 7.**
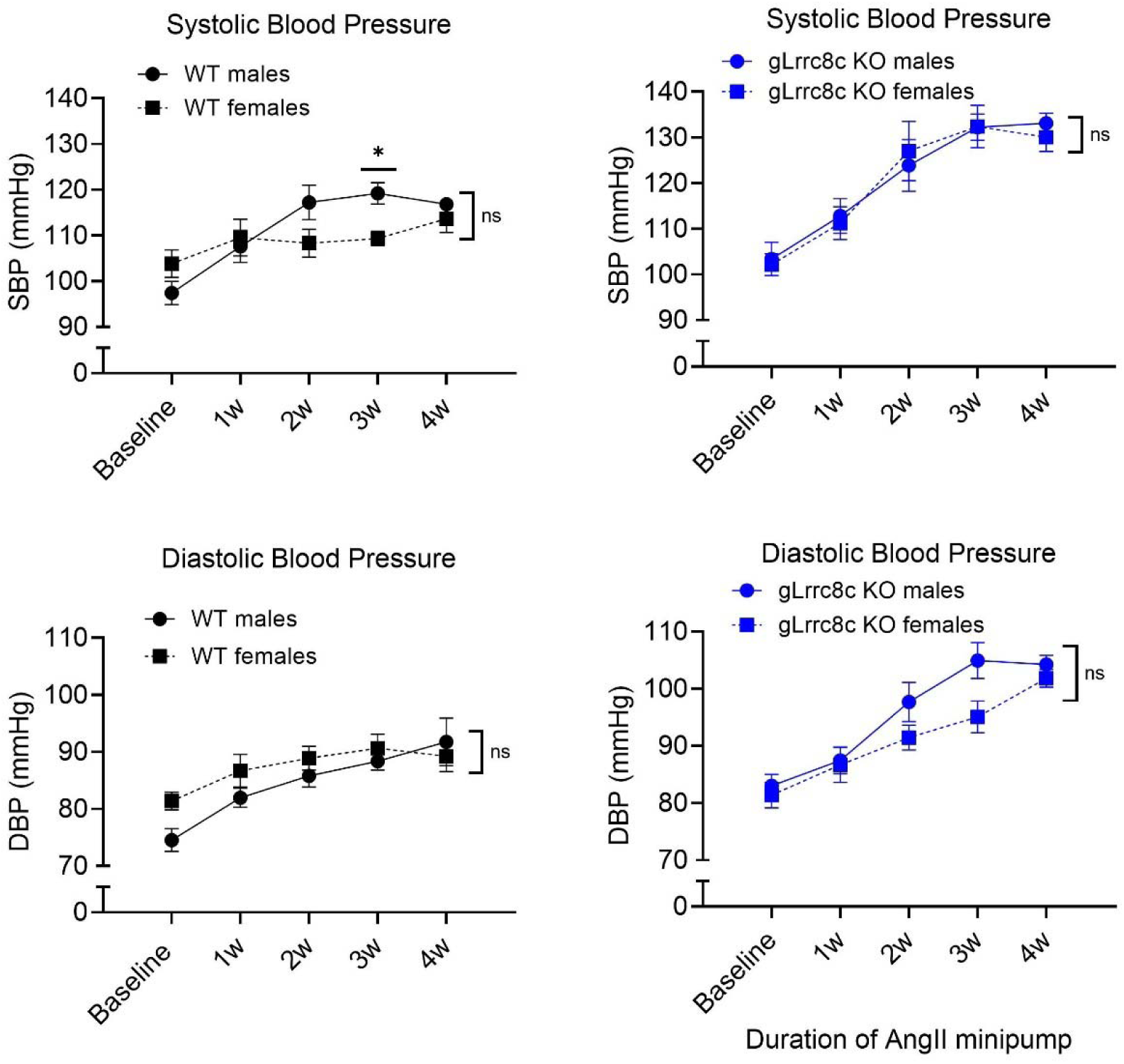
Blood pressure response to Angiotensin II infusion in male and female g*Lrrc8c* KO and control mice. Systolic **(A-B)** and diastolic **(C-D)** blood pressures in WT vs *Lrrc8c* KO mice with separated gender after 4 weeks of angiotensin infusion. Statistical significance was determined by two-way ANOVA with Bonferroni post-hoc test. Data are represented as mean ± S.E.M. ns - not significant.

### Supplemental videos

**Supplemental video 1.** *Lrrc8a*^fl/fl^ myogenic tone measured by active diameter changes of mesenteric artery during stepwise increments of pressure in the presence of Ca^2+^.

**Supplemental video 2.** e*Lrrc8a* KO myogenic tone.

**Supplemental video 3.** Wildtype control myogenic tone.

**Supplemental video 4.** g*Lrrc8c* KO myogenic tone.

**Supplemental video 5.** *Lrrc8a*^fl/fl^ mesenteric artery relaxation response to acetylcholine.

**Supplemental video 6.** e*Lrrc8a* KO mesenteric artery relaxation response to acetylcholine.

**Supplemental video 7.** Wildtype mesenteric artery relaxation response to acetylcholine.

**Supplemental video 8.** g*Lrrc8c* KO mesenteric artery relaxation response to acetylcholine.

